# Length-sensitive partitioning of *Caenorhabditis elegans* meiotic chromosomes senses proximity and number of crossover sites

**DOI:** 10.1101/2024.05.25.595906

**Authors:** Carlos Mario Rodriguez-Reza, Aya Sato-Carlton, Peter Mark Carlton

## Abstract

Sensing and control of size is critical for cellular function and survival. A striking example of size sensing occurs during meiosis in the nematode *Caenorhabditis elegans*. *C. elegans* chromosomes compare the lengths of the two chromosome “arms” demarcated by the position of their single off-center crossover, and differentially modify these arms to ensure that sister chromatid cohesion is lost specifically on the shorter arm in the first meiotic division, while the longer arm maintains cohesion until the second division. While many of the downstream steps leading to cohesion loss have been characterized, the length sensing process itself remains poorly understood. Here, we have used cytological visualization of the short arm, combined with quantitative microscopy, live imaging, and simulations, to investigate the principles underlying length-sensitive chromosome partitioning. By quantitatively analyzing short arm designation patterns on fusion chromosomes carrying multiple crossovers, we develop a model in which a short arm-determining factor originates at crossover designation sites, diffuses within the phase-separated synaptonemal complex, and accumulates within crossover-bounded chromosome segments. We demonstrate experimental support for a critical assumption of this model, that crossovers act as boundaries to diffusion within the synaptonemal complex. Further, we develop a discrete simulation based on our results that recapitulates a wide variety of observed partitioning outcomes in both wild type and previously reported mutants. Our results suggest that the concentration of a diffusible factor is used as a proxy for chromosome length, enabling the correct designation of short and long arms and proper segregation of chromosomes.

## Introduction

Sensing and control of size, position, and other geometric properties is of critical importance to diverse cellular phenomena, such as cell morphology, spindle assembly, flagellar growth, and organelle distribution. Many of these processes involve adjustment of a subcellular structure toward an ideal size and shape, as factors such as local precursor concentration and assembly rate act to equalize, for example, the lengths of two unequal flagella ^1,2^. Mechanisms such as concentration gradients, time-dependent decay, and size-proportioned attraction of cofactors may all be used as a proxy for size ^reviewed^ ^in^ ^3^. However, subcellular processes that compare the relative sizes or lengths of structures and make choices based on the result, offering a glimpse into a cell’s ability to perform computation based on size, are rarer and less well-understood. A striking example of such a process is the length sensing and subsequent functionalization of chromosome arms in meiotic prophase by the nematode *Caenorhabditis elegans*.

In *C. elegans* meiosis, a single genetic crossover occurs on each of the six chromosomes, dividing each chromosome into two “arms” of unequal length ^4–6^. The relative lengths of these arms are then sensed by an unknown mechanism, leading to the eventual recruitment, eviction, or post-translational modification of several proteins differentially on the shorter versus the longer arm. This differential functionalization of short and long arms is essential to coordinate the discrete, stepwise destruction of cohesin with the two meiotic divisions. Unlike the majority of organisms whose chromosomes each have a single centromere region, the chromosomes of *C. elegans* and most other nematodes are holocentric, with centromere activity spread along their entire lengths. Monocentric organisms use the centromere as a site to maintain sister chromatid cohesion between the first and second meiotic divisions ^7,8^. However, the lack of such a designated site in holocentric nematodes is tied to an evolutionary innovation in this process: in *C. elegans*, the short arms lose cohesion first, enabling the first meiotic division, while cohesion is maintained at the long arms until the second meiotic division. Since crossovers are generated at arbitrary positions along the chromosome, the distance between the crossover and chromosome ends needs to be sensed *de novo* on each chromosome in each meiocyte.

At the time of crossover designation, homologous chromosomes are held together by a proteinaceous structure called the synaptonemal complex (SC). The SC is made up of axial elements, which are present along the length of each homologous chromosome, and the central element, which polymerizes between homologous chromosome pairs in the process termed synapsis. The central element of the SC has been shown in *C. elegans* to exist in a phase-separated state ^9^, and is able to dynamically associate with and dissociate from chromosomes ^9–11^. The SC plays multiple roles in meiotic prophase. A primary role of synapsis is to position allelic loci on homologous chromosomes in a position favorable for proper crossover formation. In many systems, the SC has also been shown to be required for the phenomenon of crossover patterning or genetic interference ^12–15^. Recent work has shown the SC possesses a liquid crystalline structure ^9^, which is hypothesized to facilitate the rapid diffusion and coalescing of crossover-promoting factors within it ^16,17^. In *C. elegans*, the SC also plays a key role in chromosome arm length sensing and functionalization. In early prophase, both the SC central element proteins (SYP-1 through SYP-6) as well axial element proteins (the HORMA-domain proteins HTP-1 HTP-2, HTP-3, and HIM-3), localize to the entire length of synapsed chromosomes. However, concomitant with crossover designation, phosphorylation of SC central element SYP-1 by CDK-1 kinase at T452, in a polo kinase binding domain, allows the polo kinase PLK-2 to bind SYP-1; both phospho-SYP-1 and PLK-2 become enriched specifically on the short arm ^18–20^. The SC-associated meiotic RING finger proteins ZHP-1 and ZHP-2 also become enriched on the short arms at around the same time ^16^. As meiocytes transition from late pachytene to diplotene, all SYP proteins as well as ZHP-1 and −2 partition to short arms, while HTP-1 and its binding partners LAB-1 and LAB-2 partition to long arms, generating asymmetry of SC components ^5,6,19,21–24^. The partitioning of SC proteins to short and long arms are required to recruit downstream factors that lead to cohesin cleavage on the short arm, and factors that protect cohesin on the long arm, and together this enables correct chromosome segregation at meiosis I ^21,23,25^. While many steps downstream of length-sensing are known in molecular detail, the length-sensing mechanism itself remains poorly understood even at a conceptual level.

Here, we have used quantitative microscopy, live imaging, and simulations to assess the underlying rules of length-sensitive chromosome partitioning in *C. elegans* oocyte precursor cells. Normally, genetic interference limits the number of crossovers on wild-type *C. elegans* chromosomes to one ^26,27^, making the length sensing decision a simple binary choice between long and short. To help uncover the principles that govern partitioning, we took advantage of the multiple crossovers that occur on a large fusion chromosome to generate a more complex measurement problem. Based on our results, we developed a proof-of-principle simulation that can recapitulate a wide variety of partitioning outcomes both in wild-type and previously reported mutants. Our results suggest that a factor originating at crossover designation sites and accumulating within crossover-bounded, phase-separated compartments of the SC is used as a proxy for length to determine the identity of short and long arms.

## Results

### Length-sensitive functionalization of the synaptonemal complex is detectable in mid-prophase and requires phosphorylation of SYP-1

The partitioning transitions of SC elements with respect to meiosis I chromosome disjunction detailed in the Introduction are illustrated for a single chromosome in **Fig. 1A**. In a previous study, we have qualitatively shown that SYP-1 phosphorylated at T452 (hereafter, phospho-SYP-1) marks short arms ^19^. However, it remained unclear how accurately intrachromosomal length differences are discriminated to designate short and long arms, especially in the case of chromosomes with COs close to the midpoint. To investigate this question, we imaged more than one hundred wild type chromosomes to quantify the timing and extent of phospho-SYP-1 partitioning and measured the physical arm lengths in comparison. First, using immunofluorescence, we visualized phospho-SYP-1 and pan-SYP-1 (i.e., recognizing both phosphorylated and unphosphorylated SYP-1) antibody signals, as well as foci of GFP-COSA-1 that mark the sites of crossover designation (hereafter referred to as “CO sites”) in oocyte precursor cells. We imaged mid-pachytene nuclei where faint COSA-1 foci are detected and late pachytene nuclei in which 6 larger COSA-1 foci, corresponding to 6 CO sites for 6 pairs of chromosomes, are detected (**Fig. 1B**). We then traced chromosomes from each nucleus in 3D, performed computational straightening, and projected chromosomes in 2D to measure their lengths and visualize the partitioning.

**Figure 1.**
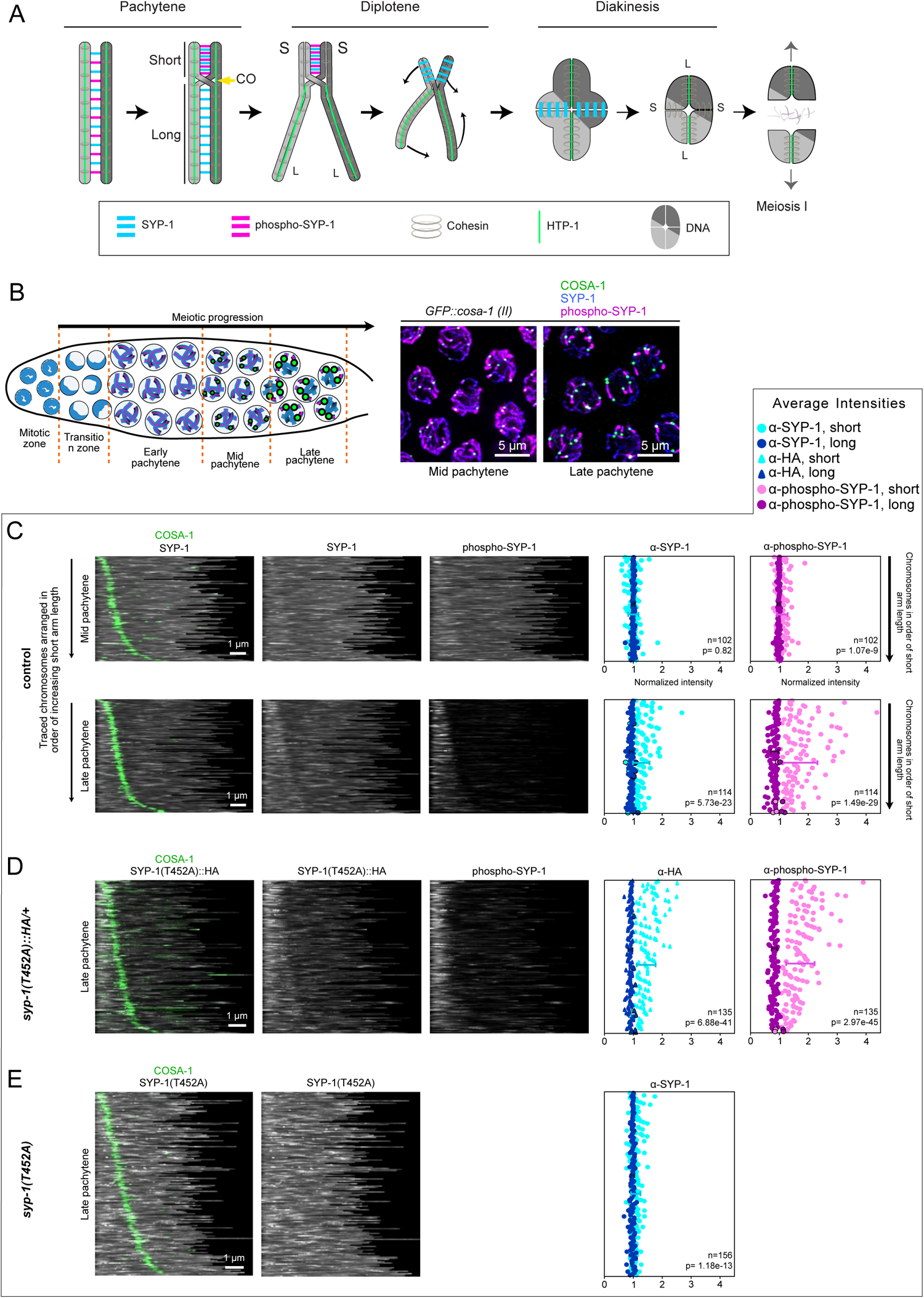
Length-sensitive functionalization of the synaptonemal complex is detectable in mid-prophase and requires phosphorylation of SYP-1. **A,** Diagram of dynamic relocalization of central (SYP-1, phospho-SYP-1) and axial (HTP-1) SC components as chromosomes remodel through late prophase. During the pachytene stage, components are initially homogeneous (left). As pachytene progresses, phosphorylated SYP-1 partitions to the short arm; by the diplotene stage HTP-1 partitions to the long arm, while all SYP-1 is found on the short arm, which becomes the site of cohesin destruction in the first meiotic division (right). **B**, *Left:* Diagram of the *C. elegans* gonad showing the timing of chromosome partitioning during prophase. The phosphorylated fraction of SYP-1 (magenta) is homogeneous throughout the chromosomes during early pachytene. During mid-pachytene, the CO sites marked with COSA-1 (green) appear as dim foci at each chromosome. At late pachytene the phosphorylated fraction of SYP-1 is restricted to the short arm region of the chromosome and the COSA-1 foci increase in size and intensity. *Right.* Representative immunofluorescence images, shown in maximum intensity projection, of mid-pachytene (before chromosome partitioning) and late pachytene (after chromosome partitioning) in *GFP::cosa-1 (II)* nuclei. The CO site protein COSA-1 (anti-GFP) is shown in green, the SC component SYP-1 (anti-SYP-1) is shown in blue, and the phospho-SYP-1 (anti-phospho-SYP-1) is shown in magenta. **C-E**. *Left*: panels show straightened images of the SC sorted from top to bottom in order of the shortest distance from the CO site (green, α-GFP::COSA-1 staining) to the nearest chromosome end and oriented such that the shorter segment is on the left. *Right:* Scatter plots of average per-arm intensity measurements of the corresponding SC images at left, sorted in the same order. The average intensity values for each segment were normalized to the average intensity of the entire chromosome and are plotted on the X axis. The blue points indicate the short segment (light blue) and the long segment (dark blue) measurements for α-SYP-1 and α-HA staining. The magenta points indicate the short segment (light magenta) and long segment (dark magenta) measurements for the α-phospho-SYP-1 staining. Outlined points are instances where the long segment average intensity is higher than the short segment average intensity. Error bars represent 1 standard deviation from the mean of the measured intensities. The p-values were obtained using Student’s t-Test comparing the mean values of the short and long segment average intensities. The n values represent the number of measured chromosomes. Panel (**C**) shows straightened *GFP::cosa-1 (II)* control chromosomes at mid pachytene and late pachytene with WT-SYP-1 (α-SYP-1) and phospho-WT-SYP-1 (α-phospho-SYP-1) staining. Panel (**D**) shows straightened chromosomes at late pachytene from *GFP::cosa-1 syp-1(T452A)/+ (II); syp-1(me17)/+ (V)* heterozygotes with SYP-1(T452A) (α-HA) and phospho-WT-SYP-1 (α-phospho-SYP-1) staining. Panel (**E**) shows straightened chromosomes at late pachytene from the non-phosphorylatable *GFP::cosa-1 (II); syp-1(T452A) (V)* mutant showing SYP-1(T452A) (stained with α-SYP-1 antibodies).

In **Fig. 1C**, straightened chromosomes are sorted from top to bottom by increasing length of the shortest segment, with shorter short segments at the top and longer short segments at the bottom. In the scatterplots at right, we plotted average intensity values of both pan-SYP-1 and phospho-SYP-1 for both short and long segments of each chromosome, normalized to the average intensity of the entire chromosome, in the same top-to-bottom order as the left panel. In mid pachytene, we detected phospho-SYP-1 signals starting to become enriched on short segments (average intensity value greater than 1) while pan-SYP-1 signals show a homogeneous distribution throughout the entire length of the SC (average intensity value close to 1). In late pachytene, we detected both phospho-SYP-1 and pan-SYP-1 restricted to the short segments, with phospho-SYP-1 more completely restricted. The mean intensity value of phospho-SYP-1 was higher for relatively small short segments and lower for relatively large short segments, suggesting that similar amounts of phospho-SYP-1 may be present on all short arms irrespective of length. Consistent with previous studies on crossover distributions ^28–30^, the majority of CO sites tend to be detected toward the ends of chromosomes. Out of 114 wild-type chromosomes traced, we observed 80 chromosomes (70%) containing a COSA-1 focus within the terminal quarters of the chromosome length adjacent to the telomeres, compared to 34 chromosomes (30%) with their COSA-1 focus located in the central half of the chromosome length. We found that in late pachytene, 110 out of 114 chromosomes had partitioned phospho-SYP-1 to the shorter segment (correctly designating physically shorter segments as functional “short arms”) while only 4 out of 114 chromosomes designated the physically longer segment as the “short arm” (*i.e.,* phospho-SYP-1 was partitioned to the physically longer segment). Of these four cases, two were the chromosomes with the highest short-to-long segment ratio (closest to 1); another contained a fairly centrally located CO site (the ratio of short segment length/long segment length is 0.77), while one had a ratio of 0.26. These exceptions show that while incorrect designation is rare overall, it may be relatively more common when arm lengths are similar. In all our observations, one side of the COSA-1 focus is always designated as the “short arm”; we found no chromosomes lacking phospho-SYP-1 partitioning. Overall, our analysis shows that chromosomes can discriminate the difference in lengths very accurately, except for rare cases of centrally located CO sites.

We next inquired about the molecular details for differential partitioning of phospho-SYP-1. Specifically, we wondered whether unphosphorylated SYP-1 completely partitions to long arms, leaving the short arm with only phospho-SYP-1 and thereby dividing the chromosome into two internally homogeneous domains, or alternatively, whether unphosphorylated SYP-1 remains distributed throughout the entire length of the SC while phospho-SYP-1 alone specifically partitions to short arms. To examine the distribution of unphosphorylated SYP-1, we placed an HA tag downstream of a non-phosphorylatable allele, *syp-1(T452A)*, and visualized SYP-1^T452A^-HA as a proxy for unphosphorylated SYP-1 (**Fig. 1D**). Surprisingly, at late pachytene in *syp-1(T452A)/+* heterozygous mutants, non-phosphorylatable SYP-1^T452A^-HA also became enriched on short arms along with wild type phospho-SYP-1 (**Fig. 1D**). Consistent with our previous results ^19^, SYP-1^T452A^ failed to partition in late pachytene in the *syp-1(T452A)* homozygous mutant (**Fig. 1E**). These observations imply that non-phosphorylatable SYP-1, and by extension unphosphorylated SYP-1, is also capable of partitioning to the short arm, but only in the presence of phospho-SYP-1.

### Phosphorylated SYP-1 restriction patterns suggest a chromosome partitioning mechanism driven by spreading and accumulation of a CO site-derived signal

To gain more insight into the underlying rules designating short and long arms, we wished to examine partitioning outcomes under multiple crossovers, which lack a simple, binary short/long arm distinction. For example, a chromosome with two CO sites, and thus three segments, would allow us to examine whether chromosome ends were a necessary part of short arm establishment, and which and how many of the three segments may become designated as the “short arm”. Wild-type *C. elegans* chromosomes implement very strong crossover interference, almost always limiting the number of crossovers on each chromosome to one ^26,27,31^. However, previous studies have shown that the triple fusion chromosome *meT7* ^27^, made by fusion of chromosomes III, X, and IV (**Fig. 2A**) and containing roughly half the genome in a single “megasome”(**Fig. S1A**), can often enjoy multiple crossovers. We reasoned that the above questions concerning mechanisms of short/long arm designation in pachytene could be unveiled by analysis of such complex cases. Further, by examining this much longer chromosome, we could ask whether any absolute size limit existed in the process of arm designation. To this end, we captured immunofluorescence images of *meT7*-containing oocyte precursor cells in late pachytene in *GFP::cosa-1 (II); meT7 (III;X;IV)* worms with 3D-SIM superresolution microscopy ^32^, which allows us to unambiguously trace the path of each chromosome, and determined the location and extent of pan-SYP-1, phospho-SYP-1, and GFP:COSA-1 (**Fig. 2B,C**). Previous studies ^27,33^ and our own analysis have shown that worms carrying *meT7* carry out meiosis with fairly high fidelity and produce viable sperm and oocytes: we found that *GFP::cosa-1 (II); meT7 (III;X;IV)* worms had an embryonic viability of 88.7% (n=1028) as compared to wild type 99.2% (n=3082). Out of 247 measured pachytene nuclei from animals homozygous for *meT7*, staining of GFP:COSA-1 at late pachytene mainly shows either 4 (54.7%) or 5 (44.1%) foci per nucleus, indicating one crossover for each normal chromosome (I, II, and V) and either 1 or 2 foci, respectively, for *meT7* (**Fig. 2B**). A small number of nuclei (3 out of 247, 1.2%) were also found with 6 COSA-1 foci, and these nuclei had 3 COSA-1 foci on the *meT7* chromosome (**Fig. S1B**).

**Figure 2.**
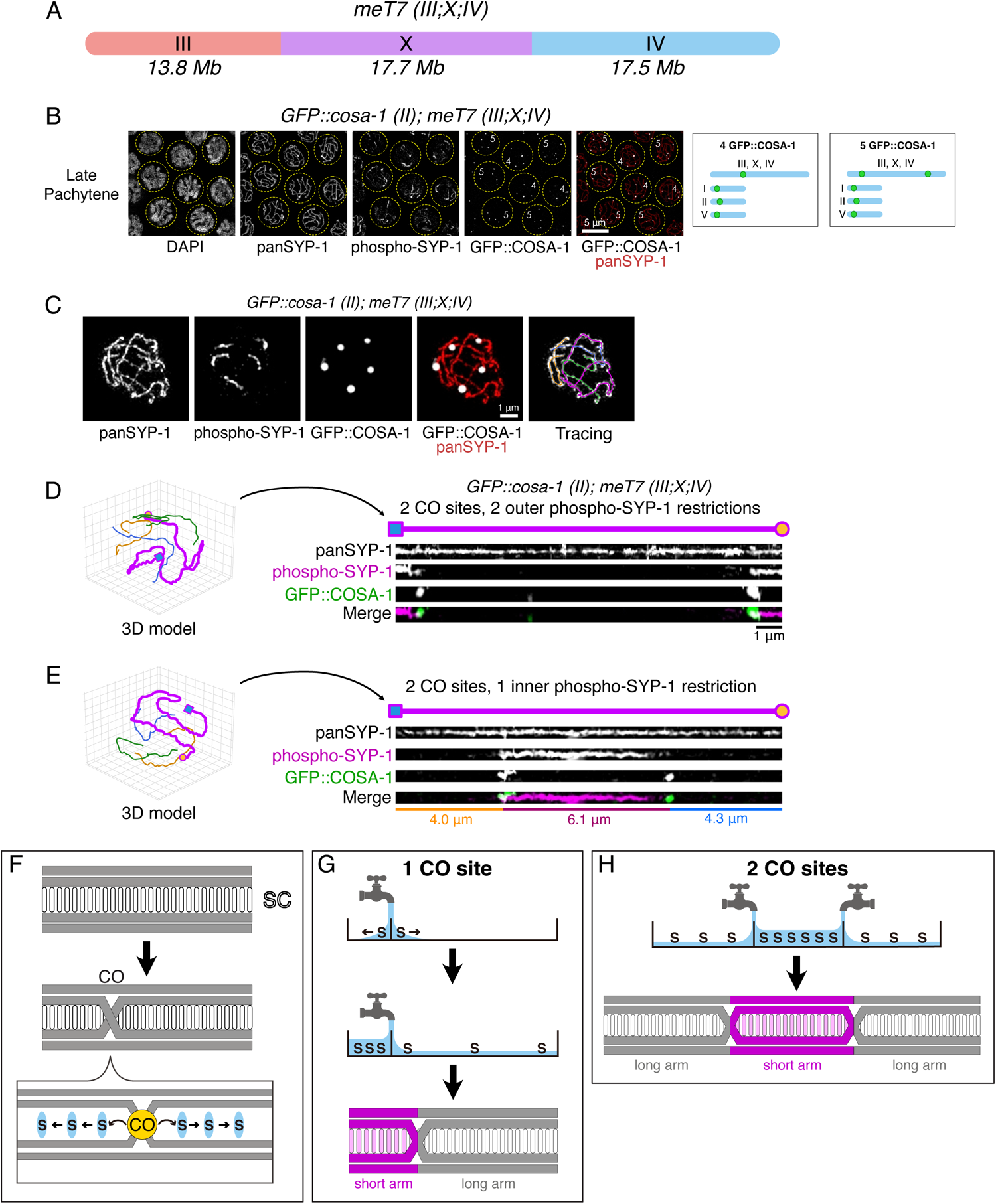
Phosphorylated SYP-1 restriction patterns suggest a chromosome partitioning mechanism driven by spreading and accumulation of a CO site-derived signal. **A.** Diagram of the triple-fusion “megasome” *meT7*, composed of *C. elegans* chromosomes III, X, and IV connected as shown. Approximate sizes (in megabases) of original chromosomes corresponding to each part of *meT7* are shown below. **B.** Visualization of CO sites in *GFP::cosa-1 (II); meT7 (III;X;IV)* marked with GFP::COSA-1 (anti-GFP) at late pachytene. The number of CO sites is shown within each circled nucleus. *Right*, karyograms indicating the distribution of CO sites per chromosome among nuclei with 4 and 5 COSA-1 foci. **C.** Representative nucleus from *GFP::cosa-1 (II); meT7 (III;X;IV)* at late pachytene showing SYP-1 (anti-SYP-1), phospho-SYP-1 (anti-phospho-SYP-1), GFP::COSA-1 (anti-GFP) and three-dimensional traces of the four chromosomes. The magenta trace represents the *meT7* fusion chromosome; blue, yellow, and green traces represent the wild type chromosomes. **D.** *Left*, Three-dimensional models of a nucleus with 5 GFP::COSA-1 foci (anti-GFP) showing the traced *meT7* in magenta and the remaining WT chromosomes in yellow, green, and blue. *Right.* straightened and maximum intensity-projected version of *meT7* showing the chromosome axis marked by SYP-1 (anti-SYP-1), two CO sites marked by GFP::COSA-1 (anti-GFP) and two outer phospho-SYP-1 (anti-phospho-SYP-1) restriction sites. **E.** As D, but for a *meT7* chromosome with 2 internal CO sites and 1 inner phospho-SYP-1 restriction site. The lengths of each chromosome segment divided by the CO sites are shown below the straightened image. **F.** *Top.* Diagram showing a synapsed chromosome before the crossover designation. *Bottom.* Diagram of the chromosome after crossover designation and a close-up on the single off-center crossover (yellow, CO) showing the diffusion of a hypothetical signal (blue, S) equally to left and right along the chromosome axis. **G.** Diagram illustrating the mechanism by which a signal originating from 1 crossover could enable length sensing of different compartments. *Top.* The hypothesized signal molecules (blue, S) are recruited or modified at the crossover and spread uniformly into two bounded unequal compartments split by the crossover. *Middle*. The accumulation of the signal leads to a higher concentration in the smaller compartment. *Bottom.* The smaller compartment with higher concentration is rectified as the “short arm” region of the chromosome. **H.** Diagram showing the expected outcome of 2 crossovers close together. *Top*. Illustration of the high-concentration inner (central) compartment in a chromosome with 2 signal origins. Flanking the inner compartment are 2 outer low-concentration compartments. *Bottom*. The high-concentration central compartment is rectified as the “short arm” region in the chromosome.

We first turned our attention to *meT7* traces with two COSA-1 foci, each localized near one chromosome end (**Fig. 2D**). At late pachytene, when pan-SYP-1 was still broadly distributed throughout the chromosome, we found phospho-SYP-1 partitioned to both COSA-1-distal segments; in other words, these chromosomes designate two “short arms” at opposite chromosome ends. This indicates that it is possible for multiple short arms to be designated on a single chromosome. In contrast, chromosomes with more centrally localized COSA-1 foci were seen with phospho-SYP-1 partitioned into the region between them (**Fig. 2E**). This observation sheds light on the mechanism of short arm determination in several ways. First, the central segment shown (6.1 μm) is longer than either distal segment (4.0 and 4.3 μm), indicating that the physically shortest segment is not necessarily the one designated as a functional “short arm”. Second, while partitioning in wild-type chromosomes always defines the short arm between a CO site and the nearest chromosome end, we find that chromosomes can also designate short arms between two COSA-1 foci. This shows that domains of phospho-SYP-1 enrichment need not contain a chromosome end or telomere sequence, and two CO sites can act as short arm domain boundaries. This is in agreement with a previous report showing an example of the *meT7* chromosome with the short arm, marked by depletion of HTP-1, detected between two chiasmata in diakinesis ^6^. Third, the central segment is roughly as long as an entire wild-type *C. elegans* chromosome, and so more than twice the length of the longest possible wild-type short arm, indicating that the typical size of short arms does not represent an intrinsic size limit to how long a “short arm” can be.

Based on these short/long arm designation observations, we developed a hypothesis that a signal is generated at or recruited to presumptive CO sites with the ability to diffuse into and within the SC (**Fig. 2F**). In wild-type chromosomes with only one CO designation site, the signal quickly accumulates to a higher density on the shorter of the two SC segments, resulting in short arm detection (**Fig. 2G**). This hypothesis makes a further specific prediction about partitioning in cases of two CO sites: since the accumulation of signal is additive, it is possible for the signal strength to be highest between the two CO sites, thus predicting short arm designation in a segment of the SC not bounded by a chromosome end (**Fig. 2H**).

### Model predictions of phospho-SYP-1 patterning in chromosomes with multiple CO sites agree with the observed meT7 chromosomes

To test the hypothesis of the short arm designation based on a diffusing signal originating from the crossovers, we developed a program that calculates the expected signal density accumulating within segments separated by crossover positions, obeying the condition that the signal is not allowed to cross from one segment to another. In the case of predictions with 2 CO sites, the signal density should be inversely proportional to the segment length, with the central segment’s density per length increased by a factor of two since it receives input from two CO sites (**Fig. 3A**). We calculated predictions based on COSA-1 focus positions from traced *meT7* chromosomes and tested whether the predictions agreed with the observed position of phospho-SYP-1 (**Fig. 3B-G**). Overall, our predictions agreed with observations of a variety of single or multiple CO cases on the *meT7* chromosome. We straightened 65 late pachytene *meT7* chromosomes with 1 GFP:COSA-1 focus and found that the crossover site tends to be designated near the center of the chromosome (41 out of 65 chromosomes had a short/long arm ratio above 0.75). The phospho-SYP-1 signal was restricted to the short segment in 54% of the cases (35 out of 65 measured megasomes) as predicted by the model (**Fig. 3B**). The remaining megasomes with 1 CO designation site showed either phospho-SYP-1 restriction in the long segment (23%) or at a small region symmetrically surrounding the CO designation site (23%) (**Fig. S1C)**. In contrast, *meT7* chromosomes bearing 2 COSA-1 foci showed a wide variety of configurations, associated with different predictions of where the putative signal would accumulate. **Figure 3** panels **C–E** show a representative set of such predictions for three possible arrangements of COSA-1 foci; in each case, the regions with the highest predicted signal level correspond to an observed region of partitioned phospho-SYP-1. The phospho-SYP-1 enrichment pattern on *meT7* megasomes with 3 CO sites also matched the regions with high signal density in the predictions (**Fig. 3F, G**). Several double- and triple-crossover chromosomes contain a central segment whose concentration is predicted to lie between that of its two adjacent segments (**Fig. 3E,F**). Interestingly, immunofluorescence images of such segments show a “mixed” partitioning in which phospho-SYP-1 is found only in a limited region directly adjacent to the COSA-1 focus. The orientation of these mixed segments was observed to follow a regular pattern: the stretch of phospho-SYP-1 starts from the COSA-1 focus that abuts the segment with lowest predicted signal concentration and extends toward the segment with highest predicted signal concentration. Taken together, these observations suggest that the short/long arm choice is made locally by comparing the concentrations to the left and right of each CO site, and segments with conflicting choices become divided between “short” and “long” identity.

**Figure 3.**
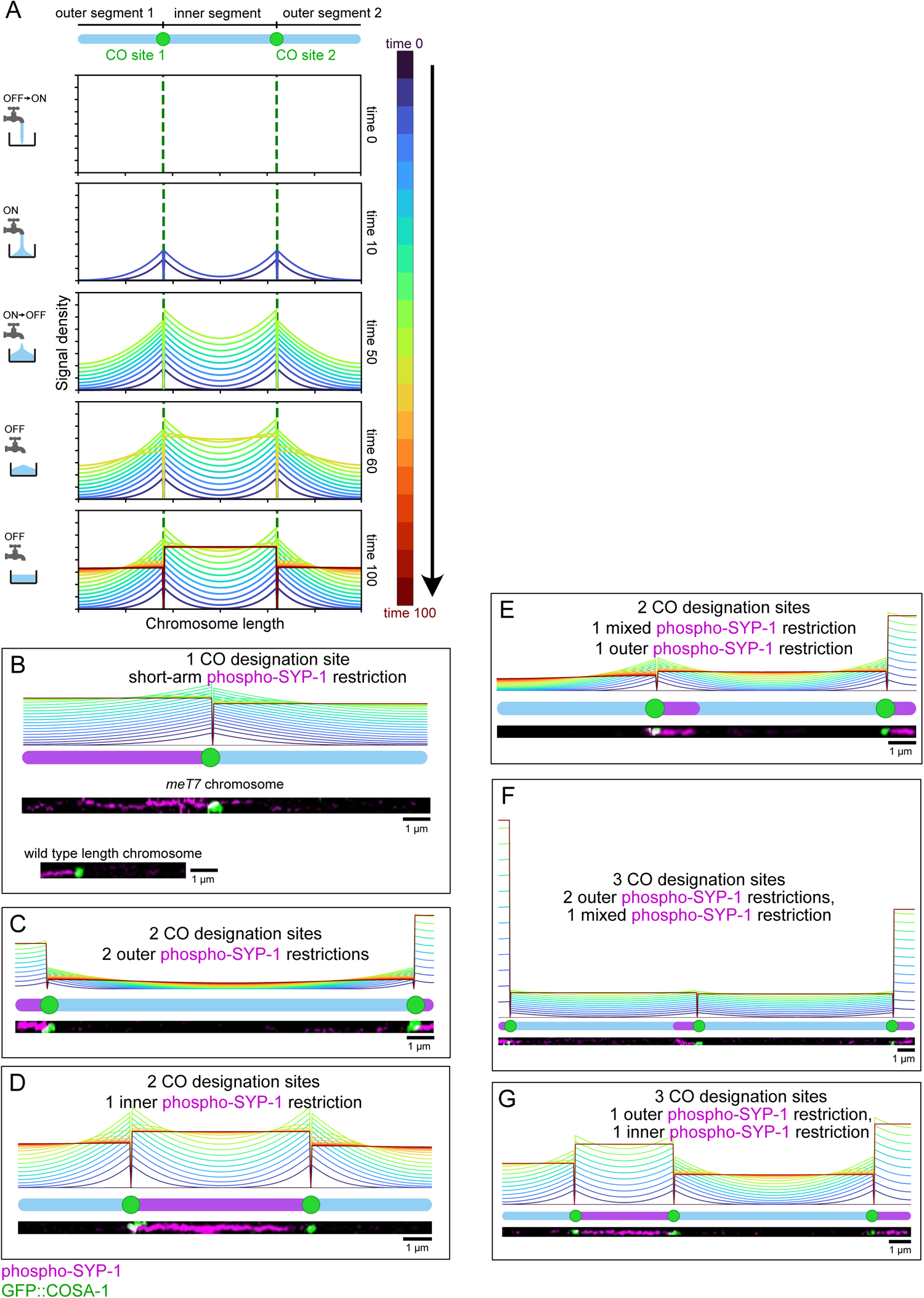
Model predictions of phospho-SYP-1 patterning in chromosomes with multiple CO sites agree with the observed meT7 chromosomes. **A**, *Top.* Diagram showing a simulated chromosome divided by two CO sites into three segments: 2 outer segments and 1 inner segment. *Bottom*. Time course of a program calculating the predictions of concentration of factors starting simultaneously at two origins. Signal density along the simulated chromosome at each time point is represented by a colored line according to the color bar shown on the left. The final signal density is shown as a red line. The green dotted lines represent the signal origin (CO sites). **B-G**, Straightened maximum intensity projection images from *GFP::cosa-1 (II); meT7 (III;X;IV)* at late pachytene with varying CO sites and positions. The images show the CO site marked by GFP::COSA-1 (anti-GFP) in green, and phospho-SYP-1 (anti-phospho-SYP-1) in magenta. The prediction of signal accumulation corresponding to the given configuration is shown above each chromosome image. Chromosome in panel D is the same as in Figure 2E.

### Quantitative analysis of two-CO meT7 partitioning agrees largely with observations

To quantitatively assess the robustness of this prediction scheme, we performed 3D-SIM imaging and tracing on 74 late pachytene *meT7* chromosomes harboring two COSA-1 foci and assessed the predicted and actual outcome of phospho-SYP-1 partitioning. To simultaneously visualize both the predicted and actual outcomes of partitioning, we plotted each chromosome as a point in a 2D space defined by the trace length-normalized position of its COSA-1 sites, arbitrarily termed CO1 and CO2. Our prediction model divides this space into three regions: distal COSA-1 sites that give rise to the lowest predicted concentration in the central segment, predicting two outer partitions (**Fig. 4A,D**); central COSA-1 sites that give rise to the highest predicted concentration in the central segment, predicting one inner partition (**Fig. 4B,D**), and mixed positions with intermediate predicted concentration in the central segment, giving one outer and one “mixed” inner partition (**Fig. 4C,D**). The zone types correspond to the final states predicted by our program for different configurations of two crossovers and are defined by where the inequalities shown in **Fig. 4D** are true. We tested the precision of the traced megasome segments lengths by retracing, straightening, and measuring the crossover positions of 3 different megasomes 9 to 10 times by 3 different people (**Fig. S2**). The resulting standard deviation of the megasome positions in the plot to their mean gives us a confidence of precision within 1% in the Y axis and 1.3% in the X axis. The predicted location and actual partitioning type of each measured chromosome is plotted in this space in **Fig. 4E**. We find that 59 out of 74 chromosomes (80%) follow the predictions as expected, i.e., the distribution of phospho-SYP-1 is constrained to the region(s) of the chromosome predicted to receive the highest levels of a crossover-derived signal.

**Figure 4.**
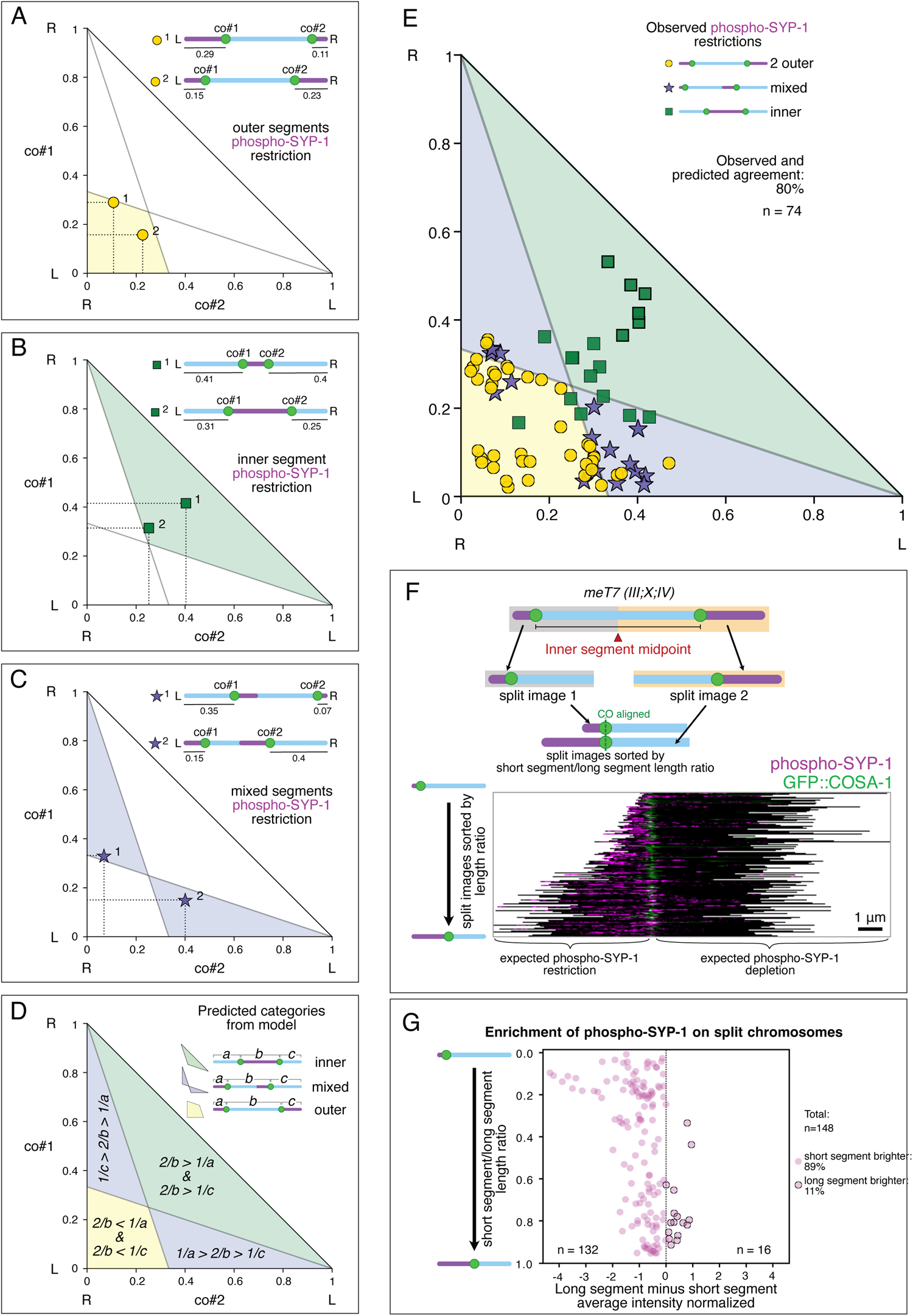
Quantitative analysis of two-CO meT7 chromosomes and model prediction accuracy. **A-C**: Phase diagrams for plotting the positions of two presumptive crossover sites marked by GFP::COSA-1 on single *meT7* chromosomes. Chromosome traces are arbitrarily assigned left and right ends and chromosome lengths are normalized to 1. The left focus (co#1) distance from the left chromosome end is plotted on the Y axis, and the right focus (co#2) distance from the right end is plotted on the X axis. The shaded region in each diagram indicates the region of the phase space in which the indicated partitioning pattern is predicted. Configurations of example chromosomes from this study with the observed GFP::COSA-1 positions are shown at upper right. **D**, Superimposition of all prediction regions, showing the distribution of each pattern type in the phase space. Each region shows equations based on segment lengths that are true within it, giving the predicted signal density at each of the chromosome segments (See **Megasome classification** in the **Detailed methods**). **E**, Plot of 74 traced chromosomes shown by markers corresponding to their observed phospho-SYP-1 restriction pattern, placed at their measured positions in the phase diagram. 80% of chromosome configurations lie within their predicted zone. Bold-outlined square markers on the top right are from measured *meT7* chromosomes whose COSA-1 foci are closer together than ⅓ of the total chromosome length. **F**, *Top.* Diagram showing division at the midpoint of two COSA-1 foci to create two half-chromosomes for individual analysis. *Bottom*. Image showing all the 148 split images coming from the megasome traces sorted by the short segment/long segment length ratio ascending from top to bottom. The split images are aligned by the GFP::COSA-1 (anti-GFP) foci. The phospho-SYP-1 (anti-phospho-SYP-1) restriction is shown in magenta. The GFP::COSA-1 channel for each split image was normalized to the corresponding GFP::COSA-1 focus area. The phospho-SYP-1 channel for each split image was normalized to the corresponding phospho-SYP-1 restriction site area. **G**, Plot showing the phospho-SYP-1 measurements of the split images from megasomes with 2 CO sites. The ratio of the short segment/long segment length is plotted on the Y axis from top to bottom. The subtraction of the long segment average intensity minus the short segment average intensity normalized to the split image average intensity is plotted on the X axis. The outlined magenta points are positive values where the long segment average intensity is higher than the short segment average intensity (16/148). The magenta points with no outline are negative values where the short segment average intensity is higher than the long segment average intensity (132/148).

While the presence of central phospho-SYP-1 partitioning demonstrates that “short arm” regions do not need to contain a chromosome end, we wondered whether residual telomere sequence from the original unfused chromosomes of *meT7* might be acting as short/long arm boundaries. We addressed this concern first by performing immunofluorescence against a FLAG-tagged version of protein TEBP-1 / DTN-1, which binds to double-stranded telomeric DNA in a sequence-specific manner ^34^, to detect whether any telomere-like structures remained at the fusion breakpoints. In cells carrying both *dtn-1::GFP::3xFLAG* and *meT7*, chromosome tracings of SYP-1 immunostaining confirmed *meT7* possessed DTN-1 foci only at the chromosome ends, suggesting that telomere structures have been lost at the chromosome fusion points (**Fig. S3A**). Next, we considered the possibility that the fusion points in *meT7* retain telomere sequence or otherwise affect partitioning without recruiting DTN-1. We simulated the effect of the *meT7* fusion points acting as boundaries for short arm formation, by plotting where partitioning would be expected to occur given random CO site positions (**Fig. S3B,C**). The resulting predictions were in striking discordance with our observations, most notably in the large number of “mixed” partitions predicted in all zones. We therefore conclude that any residual internal telomere sequence or other aspect of the fusion points on the *meT7* chromosome is unlikely to play a role in designation of partition boundaries or identity in a similar fashion to CO sites and true chromosome ends.

Although our prediction of partitioning regions based on CO position agrees well with observations, it does not predict all observed cases. We wondered whether and how often the cases where our prediction did not agree with observation for two-CO chromosomes involved a decision between two partitions of nearly equal size, similar to what we observed in the rare mispartitioning of single-CO wild-type chromosomes. To examine this quantitatively, we first simplify the analysis by dividing each two-CO chromosome case into two single-CO cases. We reasoned that if density of a factor in a segment is what determines arm identity on each side of a CO site, then bisecting the central segment (leaving density unaltered) should produce two single-CO predictions that each separately agree with the two-crossover prediction. We therefore divided each chromosome shown in **Fig. 4E** at its central segment midpoint to obtain 148 separate predictions (**Fig. 4F**) and quantitated the difference of phospho-SYP-1 intensity between the long and short segments. We found that 89% of CO sites (132 out of 148) agree with our model. Out of the 15 incorrect full length *meT7* predictions shown in **Figure 4F**, 13 had one correct and one incorrect CO site prediction, and two chromosomes had both CO sites incorrectly predicted. These instances of higher intensity of phospho-SYP-1 on the longer segment (outlined circles in **Figure 4G**) mainly occurred when the ratio of short to long segment lengths was near 1 (**Fig. 4G**). Taken together with our observations on wild type chromosome data, we conclude that the length sensing system can fail when long and short arms are very similar in length (**Fig. 1C**).

### Photoconversion imaging shows that diffusion of SYP-3 protein is impeded by CO sites

Our model for length sensing by accumulation contains the critical assumption that CO sites act as barriers that prevent or slow the diffusion of molecules within the SC, particularly the signal molecule we have hypothesized. This hypothesis is supported by the fact that COSA-1 foci mark the boundaries between short and long arms. To test this assumption experimentally, we employed *in vivo* imaging of the synaptonemal complex central element protein SYP-3 fused to the photoconvertible protein mMaple3 ^9^. We reasoned that if CO sites are barriers to diffusion, then we should expect to observe the spreading of photoconverted SC proteins on a single chromosome stopping or slowing at CO sites. To visualize COSA-1, we used a strain with a GFP:COSA-1 transgene; although the fluorescence emission spectrum of GFP overlaps with that of unconverted mMaple3, the GFP:COSA-1 foci are readily visible as large spots against the SC (**Fig. S4A**). By focusing a spot of 405 nm light on a stretch of mMaple3:SYP-3, we could achieve localized conversion of mMaple3 to the red-emitting form on one side of a COSA-1 site (**Fig. 5A**; in our figures, the red-emitting photoconverted form is shown as green on magenta to facilitate visibility). We term the side of the CO site where photoactivation was performed the “proximal” side, and the opposite side of the CO site the “distal” side (**Fig. 5A**). We visualized the spreading of the converted region using three-dimensional multiwavelength time-lapse imaging, followed by chromosome tracing, straightening, and recording of the intensity profile to test whether photo-converted SYP-3 can spread through CO sites (**Fig. 5B**). After intensity normalization and aligning all traced profiles to the center of the GFP:COSA-1 focus, we calculated the weighted average intensity of the photoactivated signal along the entire chromosome trace, and compared the COSA-1-proximal to the COSA-1-distal segments (**Fig. 5C; Fig. S4B,C**). While photobleaching reduced the overall intensity, we observed that after 9 minutes, the average intensity of the proximal (photoconverted) end was still higher than that of the distal end, indicating that photo-converted SYP-3 remained on the proximal side of the SC and did not spread through the CO site (**Fig. 5C**).

**Figure 5.**
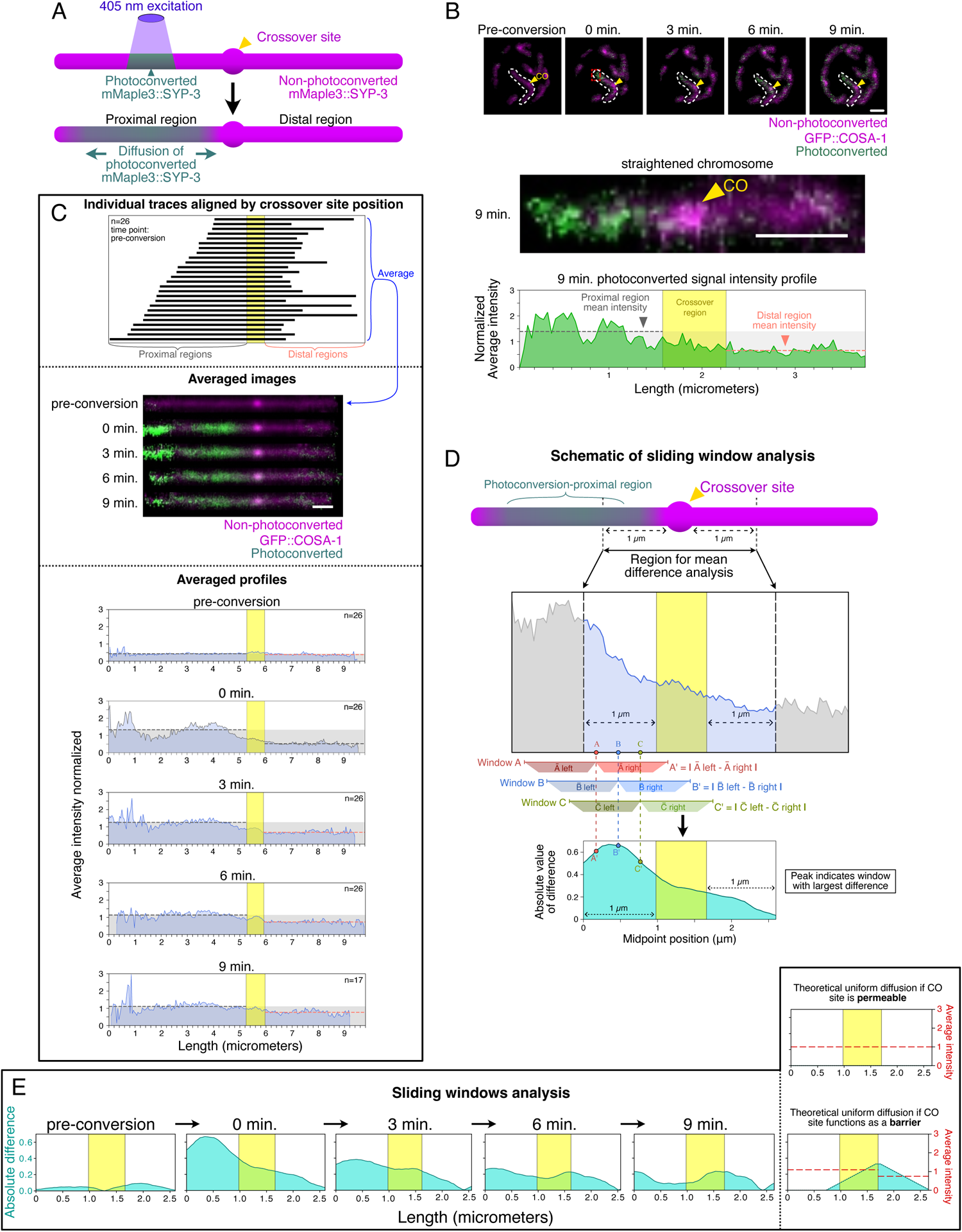
Photoconversion imaging shows that diffusion of SYP-3 proteins is impeded by CO sites. **A**, *Top.* Diagram showing the photoconversion of mMaple3::SYP-3 (shown in magenta) by 405 nm light excitation (shown in purple) to its photo-converted state (shown in green) in a limited region of a chromosome (colors intentionally chosen for visibility). The CO site is indicated with the yellow arrowhead. We term the COSA-1-demarcated region containing the photoconversion event as the proximal region, and the region past the COSA-1 focus as the distal region. *Bottom*. Diagram showing the predicted diffusion of photo-converted mMaple3::SYP-3 along the chromosome. **B**, *Top*. Merged average intensity projected images of a time course of photoconverted mMaple3::SYP-3 (green), non-photoconverted mMaple3::SYP-3 (magenta), and a GFP::COSA-1 CO site (also magenta) in one mid-pachytene nucleus of *syp-3(ok758) (I); GFP::cosa-1 mMaple3::syp-3 (II)*. The pre-conversion state is followed by 9 minutes of the time course at 3 minute intervals. Yellow arrowheads indicate the crossover position marked by GFP::COSA-1. The targeted chromosome is outlined by a white dashed line. The red box at time point *0 min.* shows the 405 nm excitation region. Scale bar: 1 μm. *Middle*. Straightened photo-converted chromosome 9 minutes after photo-conversion. Merged photoconverted mMaple3::SYP-3 (green) and non-photoconverted mMaple3::SYP-3 with GFP::COSA-1 (magenta) are shown. Yellow arrowhead indicates the crossover position. Scale bar: 1 μm. *Bottom*. Intensity profile of photoconverted mMaple3::SYP-3 from the traced chromosome 9 minutes after photo-conversion normalized to the average intensity of the entire segment. The normalized average intensity is plotted on the Y axis. The X axis represents the length of the segment in microns. The yellow rectangle is the measured extent of the GFP::COSA-1 focus region. Dotted gray and orange lines represent the mean intensity value over the lengths of the photoconversion-proximal and photoconversion-distal regions, respectively. The gray shaded region contains all values below the photoconversion-proximal region’s mean intensity. **C**, *Top.* Lineup of all 26 photoconverted straightened chromosome segments at the pre-conversion time point, shown in schematic. Segments are aligned by their GFP::COSA-1 focus midpoint. The yellow rectangle is the average crossover region of all aligned segments. The chromosome segments are oriented so that the photoconversion-proximal regions are to the left of the crossover region and sorted by the length of the region from top to bottom. *Middle.* Averaged images of all aligned chromosomes at their respective time points. The photoconverted mMaple3::SYP-3 is shown in green and the GFP::COSA-1 and non-photoconverted mMaple3::SYP-3 are shown in magenta. *Bottom.* Averaged photoconverted mMaple3::SYP-3 intensity profiles for all chromosomes at each time point. The averaged intensity values are plotted on the Y axis. The mean value over the whole length for proximal and distal regions are shown as gray and orange dotted lines, respectively. The X axis represents length in microns. **D**, *Top*. Diagram of a chromosome with the photoconverted segment (green) to the left of the CO site (yellow arrowhead) with arrows around the crossover region indicating the range used for the mean difference analysis. *Middle*. The region of the averaged photoconverted intensity profile at time point 0 used in the mean difference analysis is expanded. The blue area represents the range of the moving midpoint of the sliding window as indicated by the dashed line arrows. Three example points *A*, *B* and *C* with their respective ranges to the left and right are shown below the plot. The yellow rectangle indicates the average crossover regions of the chromosomes at 0 min. *Bottom*. Plotted values of the sliding window analysis. The absolute intensity difference of the left region from the right region surrounding each point in the range is plotted on the Y axis; the midpoint position is on the X axis. The example points *A, B*, and *C* are connected by dotted lines reaching down to their corresponding plot values, labeled *A’, B’* and *C’*. **E**, *Left.* Sliding window analysis performed on averaged chromosome traces at each time point. The absolute difference as in **D** is plotted on the Y axis. Peaks in the plot indicate points where the difference is greatest between the average intensity to the left and the average intensity to the right. *Right.* Sliding window plots of two theoretical uniform distributions of the photo-converted signal: if the crossover site was completely permeable (top) or if the right end of the crossover site functioned as a complete barrier (bottom). The values used for the sliding window analysis are plotted as red dotted lines on the Y axis. The values for the case where the crossover site functions as a complete barrier were obtained from the average intensities of the photoconverted and non-photoconverted regions at 9 min.

The differences in intensity between the proximal and distal segments could reflect a diffusion boundary at the presumptive CO site, but an alternative explanation could be that the photoactivated molecules did not have time to diffuse to the distal segment. To test whether the COSA-1 focus represents a specific intensity inflection point, we performed a sliding window analysis on each averaged time point, to highlight local changes in intensity difference within 1 μm of the COSA-1 focus (**Fig. 5D**). This analysis transforms steep intensity gradients of similar size to the window into peaks, which are more easily visualizable. We found that as time progresses, the region surrounding the COSA-1 focus shows an increasing peak in the sliding window plot, indicating a drop in average photoconverted mMaple3::SYP-3 intensity across that position (**Fig. 5D,E**). Theoretical examples of what the sliding window analysis would show if the distribution of mMaple3::SYP-3 were completely uniform within each compartment and the CO site were either completely permeable (**Fig. 5E**, right top) or completely impermeable (**Fig. 5E**, right bottom), are also shown for comparison. These results suggest that diffusion of mMaple3:SYP-3 protein is impeded across the CO site, supporting the idea that COSA-1 foci could divide the SC central element into isolated partitions.

### A simulation recapitulates partitioning of phospho-SYP-1 in wild-type and altered conditions

To explore how the local concentration of a spatially partitioned factor could in principle generate an all-or-nothing decision between segments, we implemented a discrete simulation to examine if a small number of assumptions could in principle explain the observed patterns of short/long arm partitioning. To develop the simulation, first, we assumed that a signaling factor can linearly diffuse within the phase-separated SC, as well as jump from a chromosome to an unbound pool and back to a chromosome, while CO sites act as a barrier to diffusion. Second, based on our *meT7* analysis, we assumed that the signaling factors are generated at or recruited to CO sites. Previous studies in several organisms have shown that a cyclin-like protein (CNTD1/COSA-1) and CDK-2 (cyclin dependent kinase) localize to CO sites and regulate the maturation of nascent COs, presumably by phosphorylating substrate proteins at CO sites ^20,35–39^. We imagined that phosphorylation of one or multiple substrate proteins by CDK/cyclin at CO sites might act as signals for partitioning as well. However, our model is agnostic with respect to the identities of the molecules involved, and other kinds of post-translational modifications or proteins flowing in from CO sites could serve as the signaling factor. Third, we assumed that the signal promotes its own stabilization (reducing its off-rate), thereby providing positive feedback to convert the initial asymmetry into a binary decision. We suggest this step could be mechanistically linked to the SC stabilization that occurs after CO designation, shown to consist mainly in increased SC central element binding to chromosomes but not in decreased lateral diffusion ^10,38^. Stabilization of the SC also depends on PLK-2 ^18^, particularly the PLK-1/2 dependent phosphorylation of SYP-4 at S269 ^11^. We assessed this notion by live imaging analysis, examining the spread of photoconverted mMaple3::SYP-3 in the control as well as a *syp-1* mutant (T452A), in which PLK-2 binding to the SC is prevented ^19^ ^20^. Our data showed that photoconverted mMaple3::SYP-3 spreads more quickly and to a greater extent in *syp-1(T452A)* mutants compared to the control, providing further evidence that PLK-2 slows SC dynamics through its binding to SYP-1 (**Fig. S5A,B**). Fourth, we assumed that there is a limited quantity of SC stabilizing components in the nucleus which are normally divided between six CO sites, one on each chromosome. In *dsb-2* mutants which have a reduced number of both double-strand breaks and crossovers ^40^, nuclei with 4 or more CO sites tend to partition phospho-SYP-1 correctly, while those with 3 or fewer sites usually have phospho-SYP-1 on both long and short arms ^20,41^. It has been proposed that the SC stabilization activity by PLK-2 is normally distributed across six CO sites in a wild type nucleus, but in a nucleus with fewer COs this activity will be in excess, and may spill over to the entire chromosome ^20^.

Based on the assumptions above, we implemented a simulation of discrete particles on sets of arrays, representing the SC which signal molecules can associate with (entering a specified bin of one of the SC arrays chosen at random) or dissociate from (returning to an unbound pool), allowing crosstalk to occur between different simulated chromosomes. For the wild-type case, six arrays (for the six wild-type *C. elegans* chromosomes) are simulated; for the case of *meT7,* four are simulated with one being 3× longer than the others.

In this simulation, the following actions occur at each time step (**Fig. 6A,B**): (1) signal molecules either associate (if currently unbound), dissociate (depending on a locally-determined off-rate), or randomly step left or right into an adjacent bin of the array, while not crossing specified CO positions; (2) when adjacent to a CO position, signal molecules change to an “available” state that can bind a cofactor; (3) limiting amounts of co-factor can associate randomly with available signal molecules; (4) signal molecules (bound to co-factor) recruit a stationary stabilizing factor, which decreases the local signal molecule off-rate; (5) global increase (back toward the initial, higher value) of local off-rate occurs with some probability.

**Figure 6.**
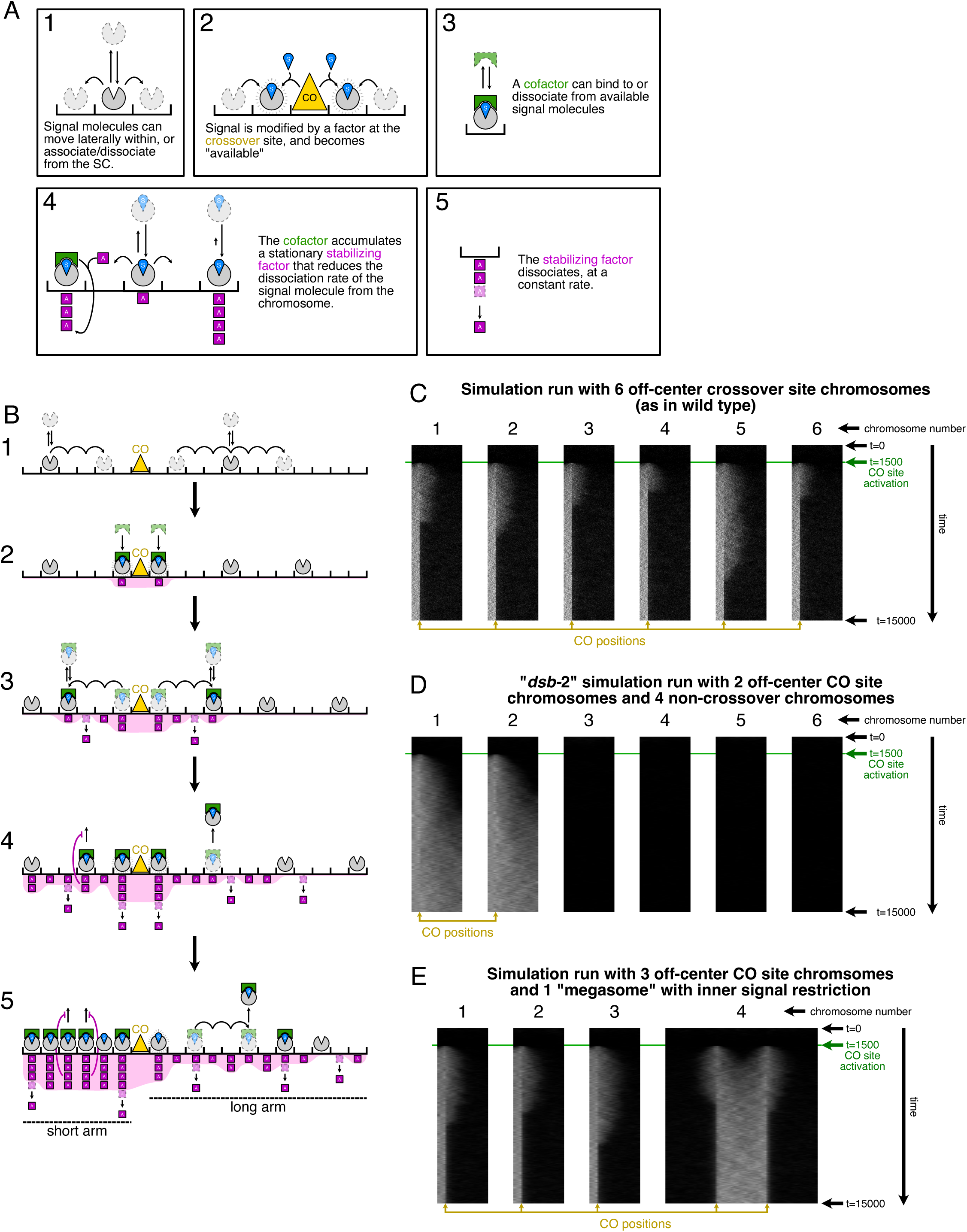
A simulation that recapitulates partitioning of phospho-SYP-1 in wild-type and altered conditions. **A**, Cartoon diagram of the 5 model steps. The simulated synaptonemal complex axis is shown as a black horizontal line with divisions to represent discrete positions along its length. 1. Signal molecules (Grey notched circles) are able to change their position in the chromosome during the simulation. The simulated molecule can move laterally within or dissociate from a simulated chromosome. Dissociated signal molecules (in a common unbound pool) can later associate with any chromosome from the entire group of simulated chromosomes at any given synaptonemal complex position. 2. Signals that move adjacent to designated crossover positions become modified and are now “available”(S, blue). 3. While available signal molecules are associated with the chromosome axis, they are able to bind molecules from a limited pool of cofactor (green). The binding is reversible. The cofactor moves with its bound signal. 4. Cofactor-bound signal molecules will deposit a stabilizing factor (“A”, magenta) at their given position. Each additional increment of stabilizing factor decreases the local probability of signal molecules dissociating from the axis at that position. The stabilizing factor does not move laterally. If signal molecules dissociate, the cofactor is released, and again becomes available to bind another chromosome-associated signal molecule. 5. The stabilizing factor is removed from the chromosome axis at a constant rate at each position. **B**, A more detailed look at events that take place as the simulation proceeds. 1. A set number of empty signal molecules at the start of the simulation will diffuse along the chromosome axis and between chromosomes. 2. Signal molecules that bump into the crossover will be modified and become available. If there are freely available cofactors, they will bind the signal molecule and commence the addition of accumulating factors at their position. 3. The cofactor bound signal will diffuse along the axis, depositing the stabilizing factor, which is also removed from the axis positions at a constant rate. 4. The stabilizing factor density will increase at a faster pace on shorter segments compared to longer segments assuming each segment has similar amounts of cofactor bound signal molecules. These shorter segments will probabilistically retain the available signal molecules longer, while the longer segments will lose them at a higher rate. 5. The outcome of this process is short segment-specific high levels of the stabilizing factor that will retain the majority of signal and cofactor molecules. **C-E,** Output examples of the simulation with distinct setup parameters. Each run consists of 15000 iterations, represented by pixel rows starting at t=0 on the top of the images and ending at t=15000 at the bottom. The X axis pixels of each chromosome represent discrete positions, with higher concentrations of the signal shown brighter. The addition of the signal starts at t=1500 when the crossover is activated (green line and label). **C**, Output of 1 run of the simulation showing the signal spreading and becoming restricted at short segments. 6 chromosomes, each divided into 100 discrete positions, were simulated. The crossover positions were set to 15 (out of 100) in all chromosomes, indicated by the yellow arrows. **D**, Output of 1 run of the simulation with 2 chromosomes with off-center crossover sites, and 4 non-crossover chromosomes. The two crossover chromosomes fail to restrict the signal to the short segments. The crossover positions were set to 15 in the two CO-containing chromosomes, indicated by the yellow arrows. **E,** Output of 1 run of the simulation with 3 chromosomes with off-center crossover sites (100 positions each) and 1 megasome (300 positions) with 2 crossover sites at positions predicted to result in inner signal restriction. The crossover positions on the normal length chromosomes were set to 15, and the crossover positions on the megasome were set to 100 and 200, indicated by yellow arrows.

The free parameters of the model include the number of signal and cofactor molecules, the initial off-rate along the chromosome, and how the off-rate increases or decreases. The chromosome lengths, crossover number, and crossover positions are also specified, allowing the simulation to be run for diverse wild-type configurations as well as altered conditions known to alter the partitioning process.

Under the above conditions, when six chromosomes with off-center CO sites are simulated, the simulation leads to positive feedback that eventually accumulates all the signal molecules on the shorter segments (**Fig. 6C**), in agreement with our observations. In developing this simulation, we found that a key aspect was making the stabilizing factor (which decreases the signal molecule off-rate) a stationary element, as opposed to the dynamically moving signal molecules. Visualizing the signal position over time as a kymograph, it can be seen that simulated signal molecules (white dots in **Fig. 6C**) begin diffusing symmetrically from crossovers, but the long arms eventually become devoid of the signal. This outcome is robustly achieved every time the simulation is run with wild type-like parameters. In contrast, when one CO site is placed near the center, the stochastic nature of the feedback often leads to errors in specification. **Figure S6A** shows five chromosomes with off-center CO sites and one with a CO site placed at 48% the distance from the left end, with correct enrichment of signal to the short arm; however, **Figure S6B** shows a summary of the end state of 100 such simulations, showing that these conditions lead to correct partitioning only 47% of the time. The other outcomes are divided between three cases: signal enrichment at the long arm (28% of the time), depletion from both arms with concomitant partial partitioning on one of the 5 wild-type chromosomes (18% of the time), and enrichment at both arms (7% of the time). This is similar to our observations on WT chromosomes with centrally placed crossovers, as we did observe partitioning of phospho-SYP-1 to the longer segment when the ratio was close to one.

Simulations in which one or more chromosomes lack crossovers entirely also recapitulated our and others’ previous observations. Just as in the *dsb-2* mutant, low numbers of CO sites in a simulation led to total failure in partitioning in most cases (**Fig. 6D, S6C**), since the excess cofactor molecules can stabilize the signal on both sides equally. With 5 CO sites, the CO-containing chromosomes all become enriched specifically on their short arms, although one of the chromosomes usually accumulates excess signal on the long arm as well, delaying its partitioning (**Fig. S6D**). Simulations of *meT7* with two CO sites also largely agreed with our imaging data; centrally positioned CO sites led to internal enrichment (**Fig. 6E, S6E**), whereas more distal CO sites led to enrichment of signal at chromosome ends (**Fig. S6F**). However, the simulation is unable to consistently reproduce the “mixed” partitioning seen in *meT7* with the same parameters; such intermediate partitions are unstable and eventually become either full or empty of the signal molecule. This outcome could indicate that one or more elements of the partitioning mechanism are not captured by the simulation, or that more fine-tuning of parameters is required. However, the basic operation of the simulation indicates that length-sensitive partitioning following the rules we have elucidated could in principle arise from molecules additively accumulating within CO-bounded compartments and changing their own dynamics in a concentration-dependent manner.

## Discussion

In this work, we have shown evidence that short arm identity, as read out by partitioning of phospho-SYP-1, is determined by the level of a factor originating from and positionally constrained by CO sites. This model leads to predictable outcomes whether single or multiple CO sites are present on a chromosome. In wild-type chromosomes, with only one CO site per chromosome, the model trivially distinguishes the physically shorter from longer arms, assuming the difference in length is large enough to generate sufficient difference in the factor levels. When multiple CO sites and thus more than two segments are present, the identity of each segment is determined additively, as a function of both segment lengths and number of bounding CO sites. We have also shown that a discrete simulation that implements positive feedback suffices to recapitulate most partitioning observations in both wild-type and several altered conditions.

While the partitioning process occasionally makes errors, in general it is capable of robustly distinguishing two chromosome segments that differ by a small amount. Typical lengths of wild-type *C. elegans* pachytene chromosomes are from 5–6μm, and cases with a short-to-long ratio of 0.75 or less are nearly all (107 out of 108) correctly decided. For a 6μm chromosome with a ratio around 0.75, differences between ∼2.6 and ∼3.4 μm segments, or a difference of 0.8 μm or larger, can be detected. This level of sensitivity is comparable to another well-characterized length-dependent process, that of *Chlamydomonas* flagellar length control, in which two flagella ∼11μm long are equalized in size when necessary and maintained at a standard deviation of ± 2.7μm ^42^. Instead of a steady-state process of size control, however, the *C. elegans* chromosome length measurement process is the precursor to a binary decision with permanent consequences for the next critical cell division of meiosis I. We found that in 3 out of the 4 cases of “incorrect” partitioning we observed in wild-type chromosomes, the arm lengths were close to 1. However, cases like the single exceptional chromosome observed whose short segment was 25% of the total length but restricted phospho-SYP-1 to the longer segment could experience problems in meiosis I disjunction, if the long arms, oriented parallel to the spindle, lost their cohesion first. However, since even mutations such as *syp-1(T452A)* that result in near-total loss of partitioning at diakinesis still have relatively high viability ^19,20^, there may be correction mechanisms redundantly ensuring correct chromosome disjunction in such cases, such as the LEM-3 nuclease that has been shown to robustly resolve chromatin bridges during chromosome segregation ^33^.

Our results on *meT7* partitioning shows that segments bounded by two CO sites have twice the “short arm generating power” per unit length than segments bounded by a CO site and a chromosome end. We suggest a parsimonious explanation of this pattern is that each CO site produces a signal that accumulates in both CO-adjacent segments, and thus acts additively between two CO sites. One possibility for such a signal is a protein that enters the chromosome at the CO site and then diffuses through the SC in either direction; another, perhaps more likely scenario is that a CO-localized activity modifies proteins already present in the SC on both sides of the CO site. While predictions based on this model are correct 96.5% of the time in the trivial case of a single CO, the success is not as high for predicting *meT7* partitioning when two CO sites are involved; out of 148 CO sites measured, only 132 (89%) were correctly predicted. Some part of this difference may be due to slight errors in tracing, as incorrect predictions often occur on the boundaries between prediction zones. Another potential source of discrepancy could be different timing of CO site “activation”, i.e., the time at which signal begins to be generated by each CO site. If one CO site on *meT7* were active long enough before the other, it could potentially change the partitioning prediction of phospho-SYP-1 from what we would predict based on the CO site locations. The errors in length measurement we observe when segments are of similar size are consistent with a previous study showing that single COs artificially introduced in the center of chromosomes fail to restrict LAB-1 to the long arm at diakinesis, leading to problems with chromosome segregation ^43^. This study also observed partitioning failure in diakinesis bivalents even with off-center COs, albeit at a lower rate; this is possibly because it was the only CO in the entire nucleus, a situation known from studies on *dsb-2* mutants with low CO numbers to cause partitioning problems at pachytene ^20,41^.

Our model of signal accumulation within segments separated by CO sites makes the assumption that the segment boundaries are mostly impermeable to the signal molecule itself. This impermeability would be necessary to create a density of signal within each segment inversely proportional to its length. Whether CO sites physically interrupt the continuity of the SC, or slow down diffusing proteins through occlusion or transient binding interactions, has yet to be conclusively determined. Observations of presumptive crossovers or recombination nodules interfacing with SC central elements have been reported in electron microscopy studies ^44,45^, and more recently in 3D-SIM fluorescence microscopy ^46^ in which SC central elements are shown to form “bubbles” surrounding COSA-1. The patterns of partitioning proteins at pachytene already strongly suggest that CO sites act as boundaries in some respect, since they tightly demarcate the short and long arms. Our live imaging experiments showing that photoconverted mMaple3:SYP-3 fails to spread through CO sites provides some evidence in support of this notion; however, since these experiments are necessarily constrained by time and photobleaching, we cannot conclusively state whether the kinetics we observe would suffice to maintain compartmentalization of signal molecules. Future experiments with more sensitive single-molecule imaging should be well-suited to address this question.

We noted several examples of closely spaced COSA-1 foci in our analysis of *meT7* chromosomes. Considering that crossover interference in *C. elegans* is known to act over the entire length of a wild-type chromosome ^13,27,35^, we expected not to observe any instances of two COSA-1 foci closer together than roughly a third of the length of *meT7* (shaded zone in **Fig. S3C**). However, we found 6 out of 74 chromosomes with COSA-1 foci spaced closer together than this length (**Fig. 4E**, dark-outlined square markers toward top right). These close foci are all quite close to the middle of *meT7*, the segment derived from the original X chromosome, suggesting that the distance between CO sites may be less on the X chromosome. One possible reason for this may be the heterochromatic nature of the X chromosome in the germline ^47^ which may reduce the length for a given number of megabases.

Our discrete simulation shows that a CO-derived signal like the one we propose could in principle cause positive feedback that partitions a signal molecule to short arms, even when the difference between arms is relatively small. The simulation also recapitulates the global partitioning failures in mutants where not every chromosome enjoys a crossover, and most of the patterns seen in *meT7* fusion chromosomes with two CO sites. However, the simulation does not stably form the “mixed” segments that are often observed by immunofluorescence on *meT7*; most simulations that start off in a pattern predicted to give rise to mixed partitioning eventually lead to signal-free zones on both sides of the CO site. The observation of mixed segments on *meT7* was unexpected, since our observations on wild-type short arms show them either full or devoid of phospho-SYP-1. The other scenario in which mixed partitions have been observed is when a single CO site occurs on *meT7*; in such cases, the zone of phospho-SYP-1 enrichment often does not reach all the way to the nearest chromosome end (see example in **Fig. S2B**). This observation was also previously made for zones of HTP-1 depletion on singly crossed-over diakinesis chromosomes of *meT7* ^6^. One possible explanation raised by our discrete simulation is that the factor does not have time to accumulate asymmetrically on one side of a relatively isolated CO site before the short arm partitions established on other chromosomes sequester all the limiting molecules for themselves. We do not know whether the mixed phospho-SYP-1 segments on *meT7* are stable or may also eventually be lost. If the actual zones of anaphase I cohesin removal were restricted to the phospho-SYP-1 regions as shown by immunofluorescence on mixed segments, it could be potential source of chromosome segregation problems, since cohesin would still hold chromatids together on both sides of a chiasma.

While the current work does not identify the factor promoting short arm identity, possible candidates are proteins lying near or at the top of the length-sensing pathway, whose mutation blocks the ability of one or more proteins to partition to the short and/or long arm. Among these are the ZHP-1/2 proteins, whose partitioning coincides with that of phospho-SYP-1 and are required for phospho-SYP-1 partitioning at pachytene ^16^. ZHP-1/2 are related to ZHP-3/4, RING finger proteins that have been shown, along with their homologs such as HEI10 in Arabidopsis, to form widely-spaced foci through a putative coarsening process at the sites of crossover designation ^17,38^. It is possible that physical parameters of coarsening could drive a similar coalescence for ZHP-1/2 at a much larger scale, such that they cover an entire chromosome segment. Similarly, minimization of surface-to-volume ratio has been proposed to aid the partitioning of the entire SC to the short arm that occurs upon desynapsis ^48^. Another candidate, not mutually exclusive with ZHP-1/2, could be PLK-2 recruited by phosphorylated SYP-1. We and others have shown that priming phosphorylation of SYP-1 at T452 by CDK-1 is required to recruit PLK-2 to the SC; PLK-2 recruitment, in turn, is required for SYP-1 partitioning ^19,20^. We have extended this observation to show that phosphorylation of SYP-1, and presumably PLK-2 recruitment, can act in *trans* to promote partitioning of a substantial fraction of non-phosphorylated SYP-1. PLK-2 has also been shown to be required for the timely partitioning of ZHP-1/2 at pachytene, but not for its later partitioning at diakinesis ^16^. PLK-2’s influence on both phospho-SYP-1 and ZHP-1/2 may be due either to a specific interaction involved in length sensing at this stage, or alternatively due to its role in inactivating CHK-2 and promoting later steps of CO designation in a timely fashion ^49^. For the former case, we hypothesize that after PLK-2 is recruited to the SC, it may phosphorylate further targets that promote partitioning by slowing down the dynamics of the SC, specifically by lowering the rate of SC dissociation from the chromosome. A candidate for such a target is S269 of SYP-4, whose phosphorylation not only affects SC dynamics but also double-strand break formation ^11^. However, partitioning of PLK-2 (and presumably of phospho-SYP-1) depends on ZHP-1/2 not only at pachytene but at later stages as well ^16^, strongly implying that ZHP-1/2 act upstream of PLK-2 and SYP-1 phosphorylation. Finally, it is possible that neither of these pathways is the initial sensor of length, but are instead responding to a prior and independent signal.

## Author contributions

Conceptualization, C.M.R., A.S. and P.M.C.; Methodology, C.M.R., A.S. and P.M.C.; Software, C.M.R. and P.M.C.; Investigation, C.M.R., A.S. and P.M.C.; Writing - Original Draft, C.M.R, A.S. and P.M.C.; Funding Acquisition, A.S. and P.M.C.

## Acknowledgments

We thank Anne Villeneuve and Ofer Rog for critical reading of the manuscript; Hiroki Shibuya, Ofer Rog and the Caenorhabditis Genetics Center, which is funded by NIH Office of Research Infrastructure Programs (P40 OD010440), for providing strains used in this study; Lexy von Diezmann and Masa Shimazoe for their advice in photoconversion experiments; Daniel Packwood for critical discussions about simulation; Sohei Sasagawa, Ryutaro Kusaba, Ryunosuke Okita, Jacky Tam, Mamoru Ishii and Ryo Takada for help with chromosome tracing; Atsushi Matsuda for 3D-SIM microscopy support; Takahiro Fujiwara and Fumiyoshi Ishidate for confocal microscopy training and support. We thank the Ministry of Education, Culture, Sports, Science and Technology and the Otsuka Toshimi Scholarship Foundation (No. 21-273) for providing scholarships. This work was supported by JSPS KAKENHI Grants #18H02373 and #22K19272 to PMC, and by the Kyoto University SPIRITS fund.

## STAR Methods text

### Resource availability

#### Lead contact

Further information and requests for resources and reagents should be directed to and will be fulfilled by the lead contact, Peter Carlton

#### Materials availability

All strains generated in this study are available upon request.

#### Data and code availability

● All data reported in this paper will be shared by the lead contact upon request.
● All original code has been deposited at https://www.github.com/carltonlab/chromosome-partitioning and is publicly available.
● Any additional information required to reanalyze the data reported in this paper is available from the lead contact upon request.

### Experimental model details

All *C. elegans* strains used were maintained at 15-25°C on nematode growth medium (NGM) and fed OP50 *E. coli* bacteria. L4 hermaphrodite worms were picked and grown for 24 hours at 20°C before live imaging and immunofluorescence experiments. The *syp-1(T452A)::HA/+* heterozygous worms used for immunofluorescence were obtained by mating *meIs8 [pie-1p::GFP::cosa-1 + unc-119(+)] II* (RRID:WB-STRAIN:WBStrain00000296) hermaphrodites with syp-1(icm81[T452A::HA]) II; syp-1(me17) V (PMC471) males.

### Method details

#### syp-1(T452A) tagging

The CRISPR/Cas9 system was used to insert a TY1::HA sequence at the C-terminus of the *syp-1(T452A) (II)* locus of the strain PMC346. A 200 bp ssDNA homologous recombination template (*c02_t452a-ha_hrt_ultramer*) was used for DNA repair and the targeting gRNA duplex was generated with the crRNA *c02_t452a_ha_crRNA2*. Adult hermaphrodites carrying the mutant allele *syp-1(T452A)* were injected at 24 hours post-L4 with the genome editing mix (17.5 µM gRNA duplex, 17.5 µM Cas9 nuclease and 6 µM ssDNA homology recombination template) to generate the *syp-1(T452A)::TY1::HA (II)* allele. The primers used during the PCR screening for the insertion candidates were *c02_t452a_ha_4_fw* and *c02_t452a_ha_4_rev*. Correct insertions were confirmed by DNA sequencing.

#### Cytology

For immunofluorescence, adult worms were picked and dissected on a coverslip in 15 µl of EBT buffer (HEPES pH7.4 27.5 mM, NaCl 129.8 mM, KCl 52.8 mM, EDTA 2.2 mM, EGTA 0.55 mM, Tween 1%, Tricaine 0.15% weight per volume, Levamisole 0.015% weight per volume). The worms were fixed by adding 15 µl of Fix solution (HEPES pH7.4 25 mM, NaCl 0.118 M, KCl 48 mM, EDTA 2 mM, EGTA 0.5 mM, Formaldehyde 0.2%) for 2 minutes. The dissected worms were sandwiched between the coverslip and a slide glass (Matsunami Inc. cat #S9901) and frozen at −80°C. The coverslip was removed using a razor blade and the slide with the worms was placed on methanol at −25°C for 1 minute. The slides were washed for 10 minutes 3 times in PBST. Slides were blocked in PBST supplemented with 0.5% bovine serum albumin for 30 minutes followed by overnight incubation of the primary antibodies in PBST at 4°C in a wet chamber to prevent desiccation. After the primary antibody incubation, the slides were washed in PBST for 10 minutes 3 times for a subsequent 2 hour long secondary antibody incubation at room temperature in a wet chamber. Following the secondary antibody incubation, the slides were washed for 10 minutes in PBST and further submerged for 10 minutes in PBST with 1 µg/ml of 4′,6-diamidino-2-phenylindole (DAPI) to stain the DNA. The slides were washed for an additional 10 minutes in PBST. Finally, a #1-S micro coverslip (Matsunami) with 13 µl of the prepared mounting medium (50 µl of 5M TRIS + 450 µl of NPG-glycerol) was used to cover the sample and it was sealed with nail polish.

#### Microscopy

For conventional fluorescence microscopy, images were acquired using a Deltavision microscope system using a 100x, 1.4NA PlanSApo objective (Olympus Inc.) with a Photometrics CoolSNAP ES2 camera (1024×1024 pixels) at a pixel size in the image plane of 64 nanometers. Raw images were subjected to constrained iterative deconvolution and subsequently corrected for chromatic aberration by sub-pixel shifting using wavelength-specific defocus values measured with polychromatic beads.

Three-dimensional structured illumination microscopy (3D-SIM) images were acquired with a Deltavision OMX Blaze microscope at the Advanced ICT Research Institute, Kobe, in the laboratory of Prof. Yasushi Hiroaka. A 60x, 1.4NA oil immersion objective was used. Wavelengths were acquired on separate cameras, processed for 3D-SIM reconstruction and then aligned to each other using brightfield images in all 4 wavelengths as a standard, using the *Chromagnon* software package ^50^.

For live imaging and photoconversion of mMaple3::SYP-3, a Zeiss LSM880 confocal microscope (Carl Zeiss) equipped with Plan-Apochromat 63x/1.40 NA objective was used. Animals were located using brightfield illumination, and nuclei suitable for image acquisition were targeted using the 488 nm laser. The excitation wavelengths for the non-photoconverted and photoconverted mMaple3::SYP-3 were 488 nm and 561 nm respectively. The software used for the image acquisition and photoconversion was *ZEN (black edition)* (Carl Zeiss) (RRID:SCR_018163).

#### Chromosome tracing, straightening and 3D model rendering

Individual nucleus images cropped from the corresponding gonad region were prepared using *Fiji* ^51^ (RRID:SCR_002285) or *Priism* ^52^ and exported to the TIFF format ^53^. The chromosome axes of the nucleus, marked by a synaptonemal complex element, were traced using a python script for *Fiji* that utilizes the *SNT* framework ^54^ (RRID:SCR_016566) or the *3dmodel* program from the UCSF Priism software suite. The chromosome traces were used to create the visual renderings of the 3D models using the *roifile* library ^55^ (RRID:SCR_023331). The traces were used to straighten the chromosomes from the original nucleus images using *Fiji*.

#### Straightened chromosome alignment and segment intensity measurement

For **Fig. 1C-E**, the crossover position for every 3D straightened chromosome was manually selected and the short segment and long segment regions were calculated using *Fiji*. The straightened chromosomes were sum projected on the Z dimension and subsequently each channel was normalized to the average intensity of the entire chromosome. The short segment region and long segment region average intensities were obtained from the normalized sum projected straightened chromosomes. To create the aligned chromosome images, the normalized sum projected straightened chromosomes were sorted by the short segment length. The vertical pixel values of the sorted chromosomes were summed at every horizontal pixel position generating 1 pixel tall images that were extended with 0 value pixels to the right to match the length of the longest chromosome of the group. The extended sorted chromosomes were combined vertically from the smallest short segment length to the biggest short segment length.

#### Model prediction

The model predictions for **Fig. 3** were calculated in python (https://www.github.com/carltonlab/chromosome-partitioning). The predicted chromosome was divided into a number of discrete units based on the length of the chromosome. The chromosome was also divided into confined segments by the specified crossover positions. The number of iterations in the prediction (represented as “time” in **Fig. 3**) was calculated based on the chromosome length to ensure diffusion throughout the chromosome. During the first half of the iterations, signal level was increased at positions directly adjacent to each crossover position, and the signal diffused further along the confined segment at a constant rate. The crossover positions stopped adding the signal at the halfway mark of all iterations, while the signal diffusion inside the confined segments continued. The resulting signal density along the predicted chromosome units at each time point was plotted and color coded according to the iteration number.

#### Megasome classification

The length of straightened megasomes with two CO sites was split into three segments delimited by the crossover sites with lengths of *a*, *b* and *c* in order from left to right. The megasomes were classified according to the pattern of phospho-SYP-1 restriction.

Megasomes with phospho-SYP-1 restricted to segments *a* and *c* were classified as *outer restriction*, megasomes with phospho-SYP-1 restricted to segment *b* were classified as *inner restriction* and megasomes with phospho-SYP-1 restriction at either segments *a* and partially to *b* or segments *c* and partially to *b* were classified as *mixed restriction*.

To determine the predicted regions by the model in the phase space diagrams of **Fig. 4A-D**, **Fig. S2** and **Fig. S3C**, the expected signal density was calculated according to the number of bounding crossovers in each of the segments. As segments *a* and *c* are only bounded by one crossover site, their expected signal density is *1*/*a* and *1*/*c*. Segment *b* is bounded by two crossover sites and thus obtains twice the signal input of segments *a* and *c* resulting in an expected signal density of *2*/*b*. The area in the phase space diagram where *1*/*a* > *2*/*b* < *1*/*c* was colored yellow and is expected to contain chromosomes with an *outer restriction* classification as the densities in segments *a* and *c* are predicted to be higher than that of segment *b*. The region described by the equation *1*/*a* < *2*/*b* > *1*/*c* was colored green and is expected to contain megasomes with a classification of *inner restriction*, where the signal density in segment *b* is expected to be higher than in segments *a* and *c*. Finally, the two remaining areas are described by the inequalities *1*/*a* > *2*/*b* > *1*/*c* and *1*/*a* < *2*/*b* < *1*/*c*, and were colored blue to represent the *mixed restriction* classification where only one of the outer segments (*a* or *c*) has a higher expected signal density than the one in segment *b*.

#### Split megasomes alignment

The straightened megasome images were split at the midpoint of the segment bounded by two crossover designation sites. The resulting images, containing one CO site each, were sum-projected and oriented so that the shorter segment is on the left. To better display the images, a manual threshold was implemented on all chromosomes to obtain the crossover designation site area and the phospho-SYP-1 restriction area and the image channels were normalized to the mean intensity of the areas. The images were then summed along the column (Y) dimension to obtain single row average images. The images were aligned by the crossover designation site and sorted by the ratio of the short segment length divided by the long segment length (**Fig. 4F**).

#### Live imaging sample preparation

*syp-3(ok758) (I); mMaple::syp-3 GFP::cosa-1 (II)* L4 staged worms were grown for 24 hours at 20°C in the dark. For imaging, 5-10 worms were picked into 15 μl of ECM buffer ^56^ (84% Leibowitz L-15 without phenol red, 9.3% fetal bovine serum, 0.01% levamisole and 2 mM EGTA) adjusted to 330 mOsm using 5M sucrose on a #1-S micro cover glass (Matsunami). The sample was then placed on a glass slide (Matsunami Inc, cat #S9901) and quickly sealed with VaLaP (1:1:1 mixture of vaseline, lanolin and paraffin) melted at 65°C.

#### Live imaging photoconversion time course acquisition

All live imaging experiments used the *ZEN (Black edition)* software (Carl Zeiss).

For **Fig. 5** and **Fig. S4**, a pre-conversion 3D image stack from the mid pachytene region of *syp-3(ok758) (I); GFP::cosa-1 mMaple3::syp-3 (II)* gonads was acquired and used to select 2-3 chromosomes with 1 distinguishable GFP::COSA-1 focus that share the same z-position. A small region at the desired distance to the crossover was drawn on top of the non-photoconverted mMaple3::SYP-3 signal for each of the selected chromosomes.

Photoconversion of mMaple3::SYP-3 at the drawn regions was achieved using 35% power of the 405 nm laser in a single iteration with a speed parameter of 6, resulting in a pixel dwell time of 3.98 µs. Following the photo-conversion, a total of 4 3D-images at 3 minute intervals were obtained in a time course starting at t=0 min. and ending at t=9 min.

Between each time point acquisition, the stage x, y and z coordinates were manually adjusted to compensate for the nuclear and gonad movements. The zoom parameter for the image acquisitions was set to 6 with dimensions of 528×528 resulting in a pixel size of 42.6 nm and spacing of 0.35 μm in the z-dimension. The speed parameter for the image acquisition was also set to 6, resulting in a pixel dwell time of 3.98 μs.

For **Fig. S5**, nuclei from the late pachytene region of *syp-3(ok758) (I); GFP::cosa-1 mMaple3::syp-3 (II)* and *syp-3(ok758) (I); GFP::cosa-1 mMaple3::syp-3 (II); syp-1(T452A) (V)* were selected for photoconversion. A pre-conversion 3D image stack was obtained followed by photoconversion of 25-50% of the area from 3-4 nuclei from the gonad region. Images were taken with a zoom parameter of 6 and image dimensions of 512×512.

Photoconversion was achieved with the 405 nm laser power set to 45% with a speed parameter of 6. 5 time points were taken after photo-conversion with 5 minute intervals starting at *t=0min* and ending at *t=20 min*. The stage x, y and z coordinates were manually adjusted to compensate for the nuclear and gonad movements.

#### Photoconversion time course background subtraction

The photoconverted and non-photoconverted channels from the photoconversion time course images in **Fig. 5** obtained with the Zeiss LSM880 confocal microscope were separated and saved in TIFF format for further processing. First, a 3D mask of the chromosome axes and the crossover sites from the non-photoconverted channel was generated using *Fiji* ^51^ and the *Trainable Weka Segmentation* plugin ^57^. The non-photoconverted mask was used to obtain an image without the pixels outside of the chromosome axes and crossover designation sites referred to as the *masked non-photoconverted image*. The non-photoconverted mask was later used to obtain the background signal corresponding to all the pixels outside the chromosome axes and crossover designation sites. The mean intensity of every z-section of the background was measured with *Fiji* and subtracted from the corresponding z-section of the *masked non-photoconverted image* to obtain the final background subtracted non-photoconverted image. To subtract the background signal from the photoconverted channel, the non-photoconverted mask was used to generate an image of the photoconverted channel with only the pixels inside of the chromosome axes and crossover designation sites referred to as the *masked photoconverted image*. The background signal for the photoconverted channel was obtained by removing the pixels corresponding to the photoconverted nuclei. The mean intensity of every z-section of the photoconverted background was measured and subtracted from the corresponding z-section of the *masked photoconverted image* to obtain the background subtracted photoconverted image.

#### Diffusion measurements of photoconverted mMaple3::SYP-3

All 98 nucleus from **Fig. S5C** from *syp-3(ok758) (I); GFP::cosa-1 mMaple3::syp-3 (II)* and *syp- 3(ok758) (I); GFP::cosa-1 mMaple3::syp-3 (II); syp-1(T452A) (V)* were pooled together and manually classified according to the percentage of the nonphotoconverted signal where the photoconverted signal had diffused. The categories were 0%, 1-25%, 26-50%, 51-75% and 76-100%. The filenames of the images for both genotypes were obscured and the order was randomized for blind classification.

## Quantification and statistical analysis

### Quantification and statistical analysis of straightened chromosome segment average intensities

Descriptions of the statistical analysis and group sizes for the short and long segments measurements in **Fig. 1C-E** are included in the figure legend and details about the segment intensities measurement can be found in the ***Straightened chromosome alignment and segment intensity measurement*** section of the **Method details**. The n values represent the number of straightened chromosomes measured. The python library SciPy ^58^ (RRID:SCR_008058) was used to obtain the standard deviations and to compare the means of the short segments and the long segments average intensities using Student’s t-Test.

### Quantification of phospho-SYP-1 enrichment in split meT7 chromosomes

Description of the group sizes of the quantification of phospho-SYP-1 enrichment in the split *meT7* chromosomes can be found in the legend of **Fig. 4**. The short segment average intensity and the long segment average intensity of every split chromosome was measured and the subtraction of the long segment average intensity minus the short segment average intensity was plotted in **Fig. 4G**.

## Supplemental information titles

**Document S1.** Figures S1-S6. Related to Figures 1-6.

**Video S1.** Predicted signal patterning for chromosomes with multiple crossover sites. Related to Figure 3.

Predictions showing the signal density in 6 different cases of chromosomes with varying crossover positions and numbers. The colored lines indicate signal density and time point according to the color bar at the right. *Top left.* Case with 1 crossover position. *Top center*. Case with two crossover resulting in 2 outer signal restrictions. *Top right.* Case with two crossovers resulting in one inner signal restriction. *Bottom left.* Case with 2 crossovers resulting in 1 outer and 1 mixed signal restrictions. *Bottom center*. Case with 3 crossovers resulting in 2 outer restrictions and 1 mixed restriction. *Bottom right.* Case of 3 crossovers resulting in 1 outer and 1 inner restrictions.

**Video S2.** Photoconverted mMaple3::SYP-3 diffusion is restricted by the crossover position. Related to Figure 5.

Time course showing the diffusion of photoconverted mMaple3::SYP-3 (shown in green) in a *syp-3(ok758) (I); GFP::cosa-1 mMaple3::syp-3 (II)* nucleus. The video shows non-photoconverted mMaple3::SYP-3 and GFP::COSA-1 in magenta. The time course shows the nucleus before photoconversion (pre-conversion) and four succeeding time points with 3 minute intervals starting *0 min*. and ending at *9 min*. 405 nm light was used to photoconvert mMaple3::SYP-3 at a small chromosome region shown at *0 min.* (red box).

**Video S3.** Four different simulation runs of chromosome groups with diverse conditions. Related to Figure 6.

Video showing the output of the simulation of signal diffusion in 4 different chromosome groups. The simulated signal molecules are added at the crossover sites and diffuse throughout the chromosomes according to the model (See Figure 6). A total of 15000 iterations are shown starting at t=0 and ending at t=15000. The crossovers are activated at t=1500. The windows show 50 iterations at once. The normal length of chromosomes is set to 1 while the megasome length is set to 3.

A. Group of 6 normal length chromosomes with one off-center crossover position each set at 0.15.

B. Case with 2 normal length chromosomes with off-center crossovers set at 0.15 and 4 normal length non-crossover chromosomes.

C. Case with 3 normal length chromosomes with off-center crossovers set at 0.15 and 1 megasome with two crossovers set at 1.0 and 2.0 resulting in inner restriction of the signal.

D. Case with 3 normal length chromosomes with off-center crossovers set at 0.15 and 1 megasome with two crossovers set at 0.5 and 2.5 resulting in outer restriction of the signal.

