## Supplemental Data and Methods for "Length-sensitive partitioning of *Caenorhabditis elegans* meiotic chromosomes senses proximity and number of crossover sites"

### Figure S1

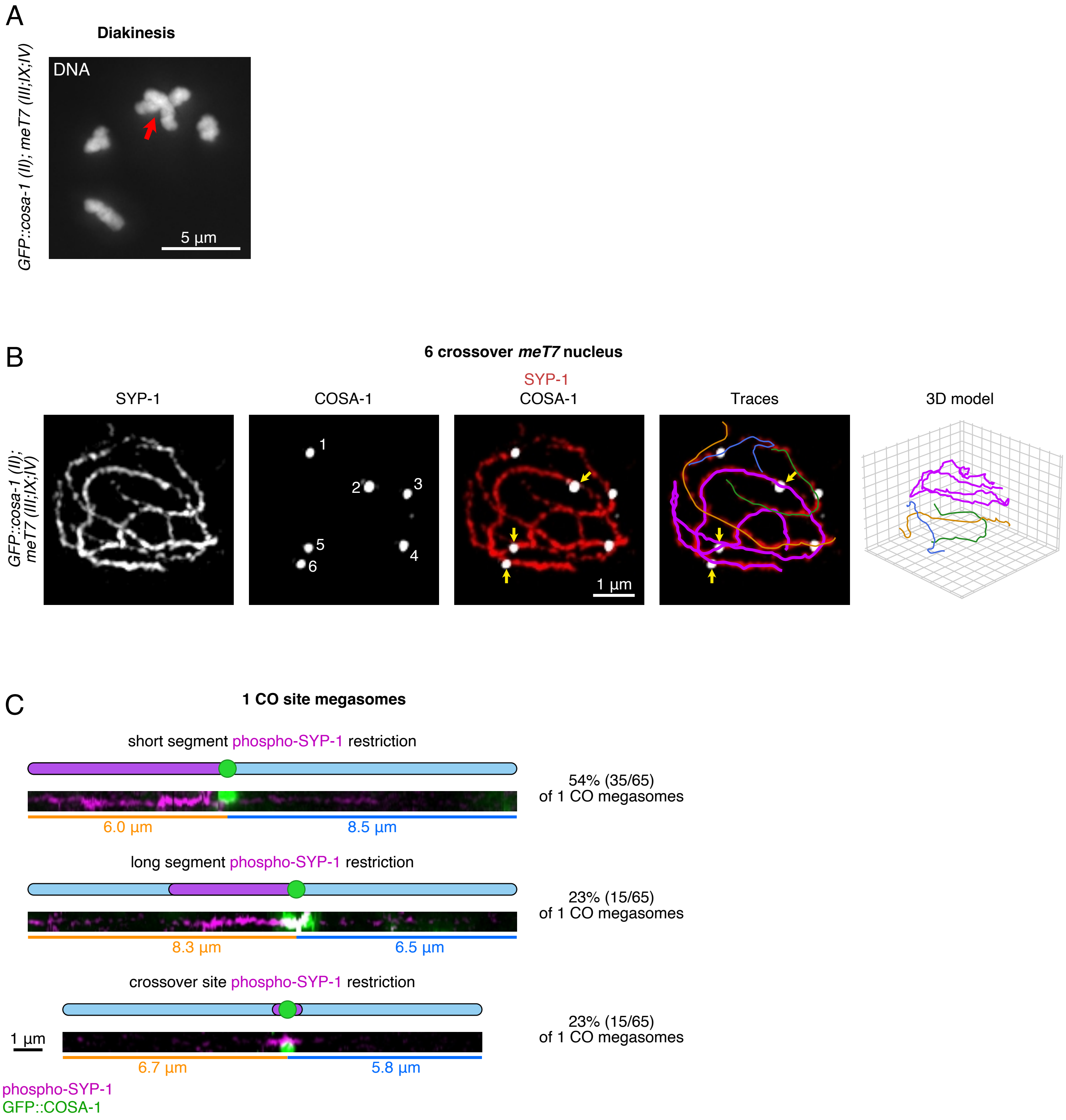
A
Diakinesis
GFP::cosa-1 (II); meT7 (III;IX;IV)
DNA
5 µm
B
6 crossover meT7 nucleus
GFP::cosa-1 (II); meT7 (III;IX;IV)
SYP-1
COSA-1
1
2
3
4
5
6
SYP-1
COSA-1
1 µm
Traces
3D model
C
1 CO site megasomes
short segment phospho-SYP-1 restriction
6.0 µm
8.5 µm
54% (35/65)
of 1 CO megasomes
long segment phospho-SYP-1 restriction
8.3 µm
6.5 µm
23% (15/65)
of 1 CO megasomes
crossover site phospho-SYP-1 restriction
1 µm
6.7 µm
5.8 µm
23% (15/65)
of 1 CO megasomes
phospho-SYP-1
GFP::COSA-1

**Figure S1.** *Single and triple crossover designation sites in meT7*. Related to Figures 2-3.

**A**, DAPI stained nucleus at the diakinesis stage in the strain *GFP::cosa-1 (II); meT7 (III;X;IV)*. Four DAPI bodies are observed. Red arrow indicates the large *meT7* triple-fusion chromosome.
**B**, *meT7* traces in a late pachytene nucleus with 6 detected GFP::COSA-1 CO sites. Images show SYP-1 (α-SYP-1) as the chromosome axis marker and GFP::COSA-1 (α-GFP) as the CO site marker. The yellow arrows indicate the 3 GFP::COSA-1 foci on the megasome.
**C**, Straightened *meT7* megasomes with 1 GFP::COSA-1 focus (α-GFP) showing phospho-SYP-1 (α-phospho-SYP-1) restriction. *Top*. Megasome with phospho-SYP-1 restricted to the short segment of the chromosome. *Middle*. Megasome with phospho-SYP-1 restricted to the long segment of the chromosome. *Bottom*. Megasome with phospho-SYP-1 restricted to the crossover designation site of the megasome.

### Figure S2

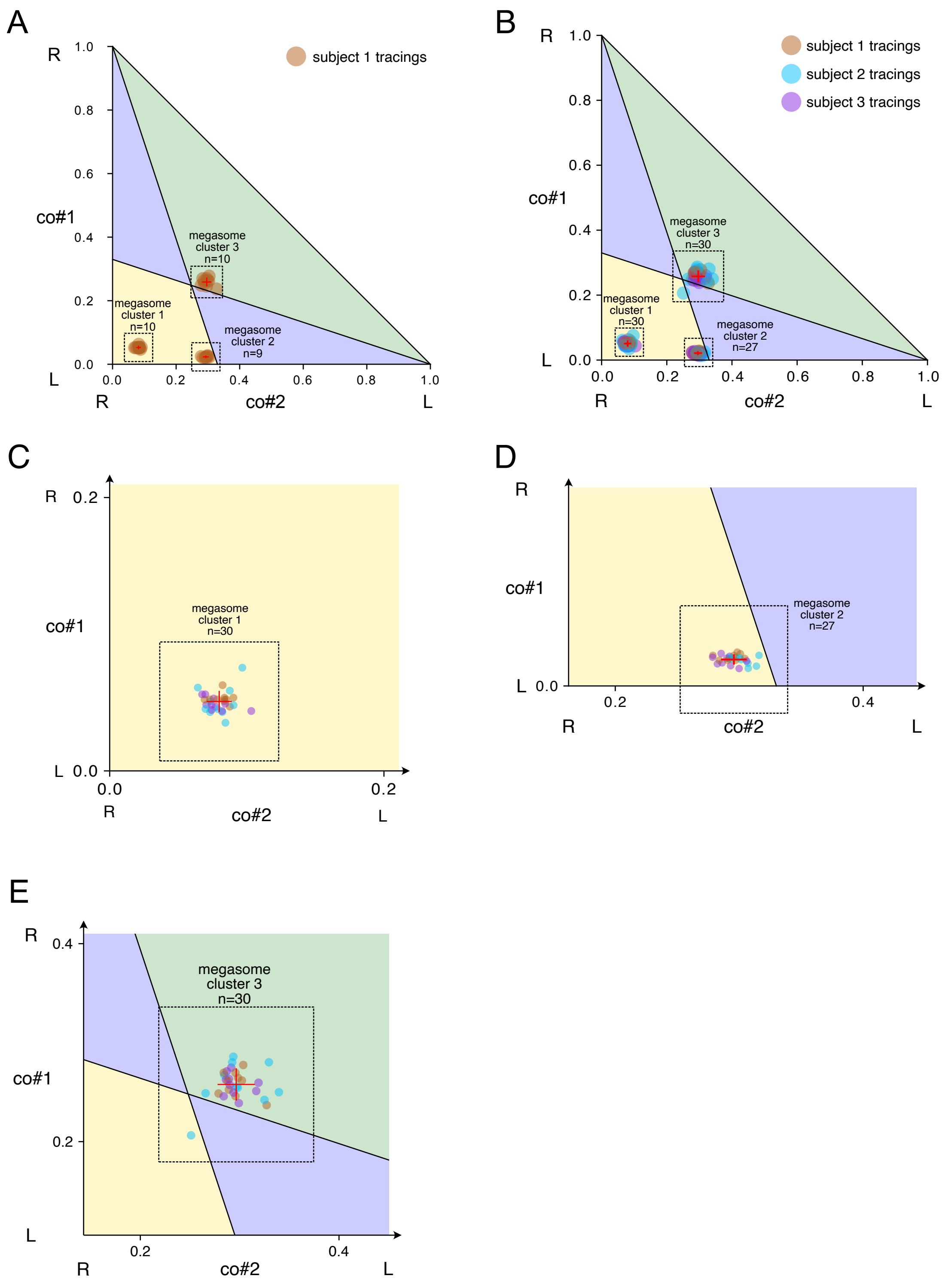

A
subject 1 tracings
megasome cluster 1 n=10
megasome cluster 2 n=9
megasome cluster 3 n=10
co#1
co#2
R
L
B
subject 1 tracings
subject 2 tracings
subject 3 tracings
megasome cluster 1 n=30
megasome cluster 2 n=27
megasome cluster 3 n=30
C
megasome cluster 1 n=30
D
megasome cluster 2 n=27
E
megasome cluster 3 n=30

**Figure S2**. *Precision of megasome tracing, straightening and CO site position measurements.* Related to Figure 4.

**A-B**, Phase diagrams showing the results of repeated blind tracings, straightening and GFP::COSA-1 focus position measuring for three different *meT7* chromosomes. All three were randomly presented 10 times each, in different rotations and/or reflections. Three clusters areplotted, each corresponding to a different chromosome. Red bars represent 1 standard deviation from the mean of the cluster.

**A**, Phase diagram for the tracings by subject 1 alone.

**B**, Phase diagram for the tracings by 3 different subjects, color-coded by subject.

**C-E**, Magnification of the phase diagram over the megasome clusters to show the spread of megasome tracings with more detail.

**C**, Magnification of the subset cluster 1 (n=30).

**D**, Magnification of the subset cluster 2 (n=27).

**E**, Magnification of the subset cluster 3 (n=30).

### Figure S3

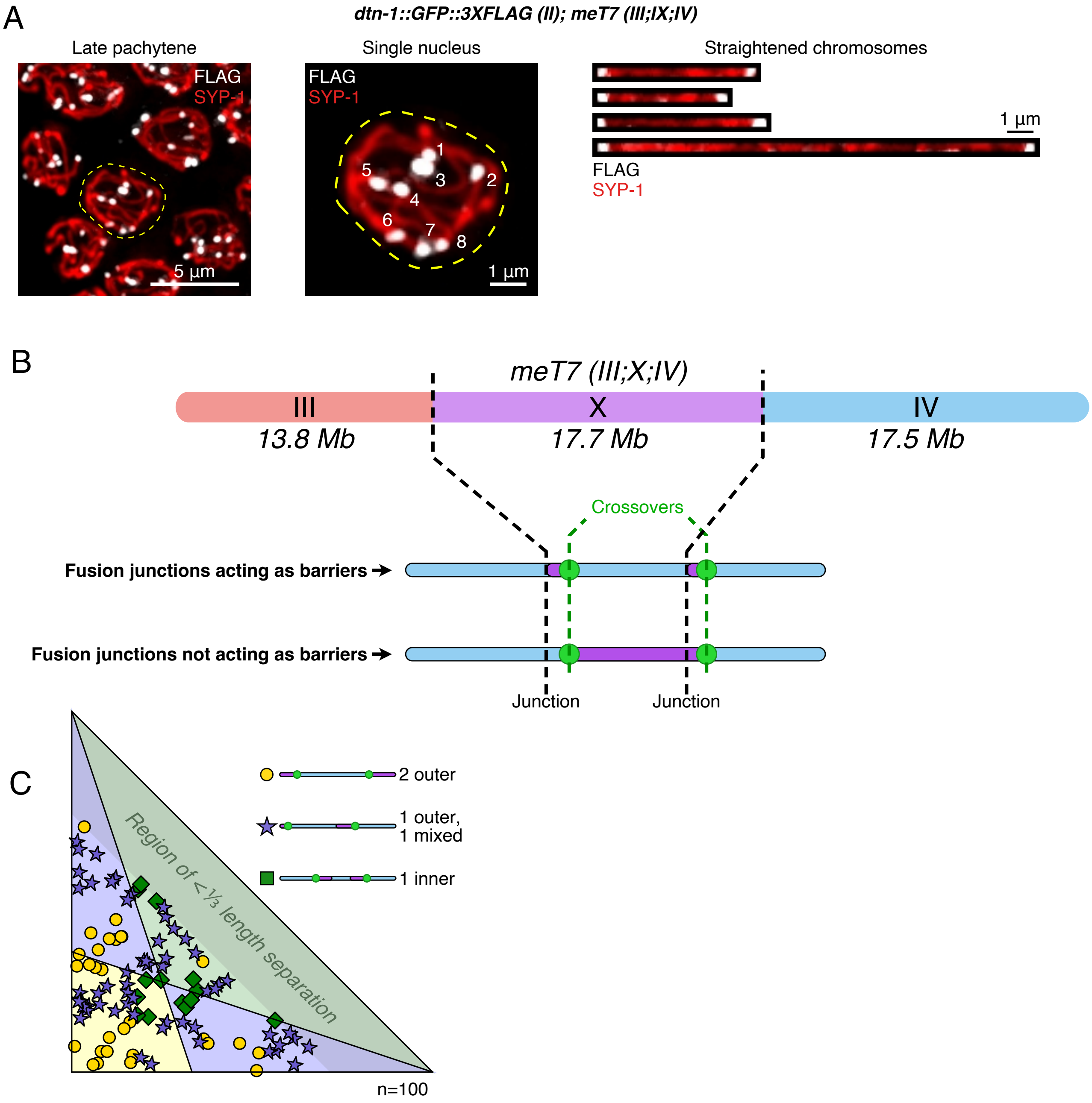


Predictions for *meT7* fusion points as barriers

**Figure S3**. *Inner junctions of meT7 do not explain the observed phospho-SYP-1 patterning*. Related to Figures 3-4.

**A**, Immunofluorescence staining of the telomere binding protein DTN-1 (white, anti-FLAG) in nuclei of *dtn-1::GFP::3XFLAG (II); meT7 (III;X;IV)* animals. The SC protein SYP-1 (red, anti-SYP-1) was used as the chromosome axis marker. *Left.* Representative late pachytene region showing DTN-1 at the telomeres. *Center*. Magnification of the outlined nucleus showing 8 foci of DTN-1. Nearby nuclei in the region were cropped out of the image (replaced with black) for clarity. *Right*. Straightened chromosomes of the outlined nucleus showing two DTN-1 foci per chromosome.

**B,** *Top.* Diagram of the *meT7* chromosome with highlighted junctions (black dotted lines) between the fused chromosomes. The approximate fused chromosome sizes are denoted in megabases. *Bottom.* Two chromosome diagrams with the same length, crossover position and junctions showing different expected phospho-SYP-1 restriction patterns. The top chromosome results in 1 outer segment and 1 mixed segment if the junctions behave as barriers. The bottom chromosome shows a single, full inner segment restriction if the junctions do not behave as barriers, as predicted by the model (See **Fig. 3**).

**C**, Phase diagram containing a total of 100 simulated megasomes with two randomly placed crossovers with the minimum distance between them set to the approximate length of chromosome III. The phospho-SYP-1 restriction prediction was generated with the assumption that the junctions do function as barriers and plotted as yellow circles (2 outer restrictions), purple stars (mixed restriction) and green boxes (1 inner restriction). The regions in the phase space diagram are calculated as in **Fig. 4**, according to the prediction where the junctions do not function as barriers.

### Figure S4

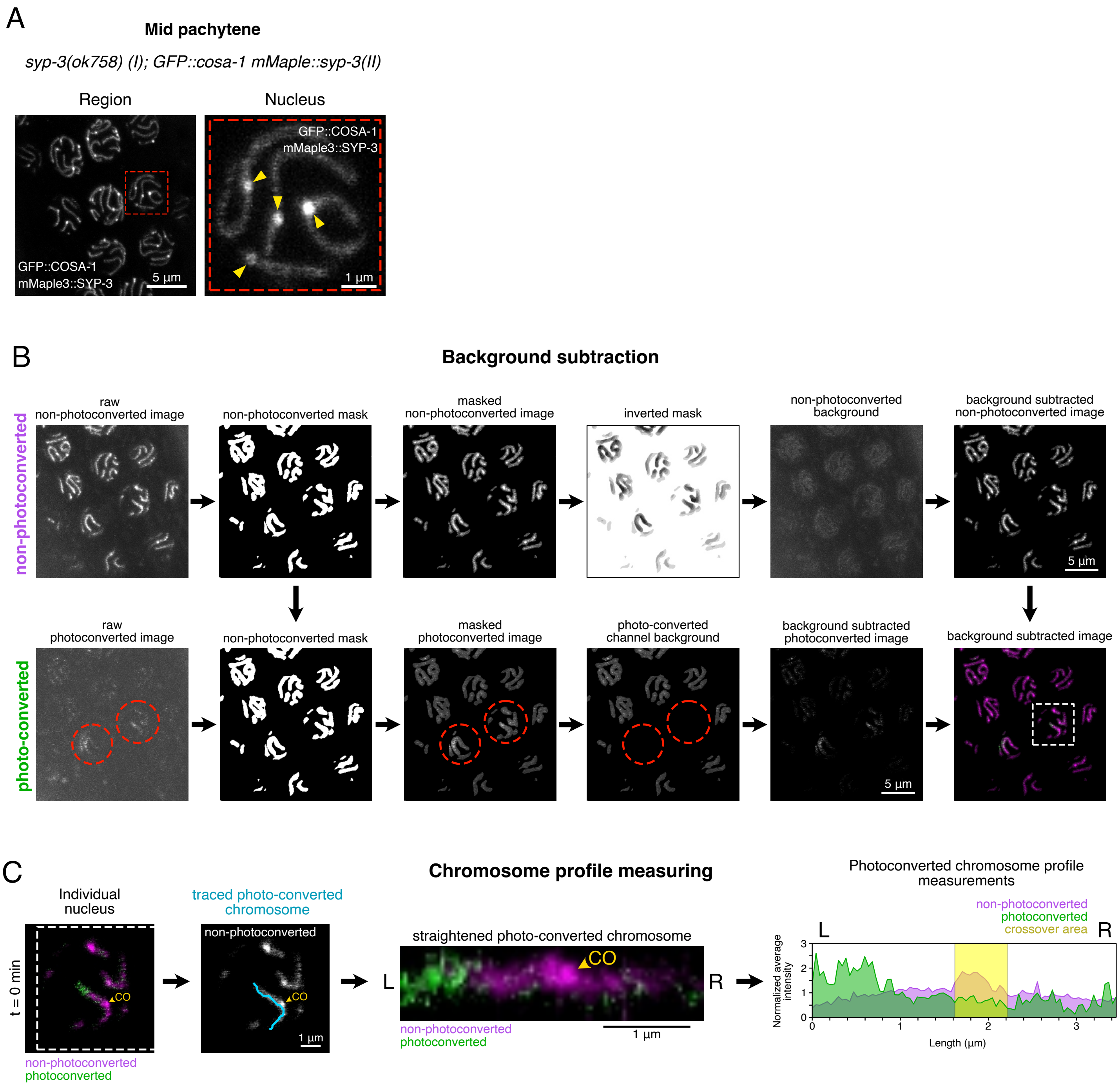

A
Mid pachytene
syp-3(ok758) (I); GFP::cosa-1 mMaple::syp-3(II)
Region
Nucleus
GFP::COSA-1
mMaple3::SYP-3
5 µm
1 µm
B
Background subtraction
non-photoconverted
raw
non-photoconverted image
non-photoconverted mask
masked
non-photoconverted image
inverted mask
non-photoconverted
background
background subtracted
non-photoconverted image
photo-converted
raw
photoconverted image
masked
photoconverted image
photo-converted
channel background
background subtracted
photoconverted image
background subtracted image
C
Chromosome profile measuring
Individual
nucleus
t = 0 min
CO
non-photoconverted
photoconverted
traced photo-converted
chromosome
straightened photo-converted chromosome
L
R
Photoconverted chromosome profile
measurements
crossover area
Normalized average
intensity
Length (µm)

**Figure S4**. *Non-photoconverted and photoconverted mMaple::SYP-3 signal background subtraction and straightened chromosome profile measurement*. Related to Figure 5.

**A**, Mid pachytene region and single nucleus of *syp-3(ok758) (I); GFP::cosa-1 mMaple3::SYP-3 (II)* showing the overlapping fluorescent emission of GFP::COSA-1 and mMaple3::SYP-3. *Left.* Max intensity gonad region projection. *Right.* Close up of a nucleus from the gonad region on the left. Yellow arrows indicate the CO sites marked by GFP::COSA-1 clearly distinguishable from the mMaple3::SYP-3 marked chromosome axis.

**B**, Background subtraction steps for the photoconverted and non-photo converted mMaple3::SYP-3 channels in the *syp-3(ok758) (I); mMaple3::syp-3 (II)* strain. The images shown are max intensity projections. *Top.* The *raw non-photoconverted image* is used to obtain a *non-photoconverted mask* from the chromosome axes. The mask is employed to create the *masked non-photoconverted image* that lacks numerical values (pixels set to "NA") outside the chromosome axes region. An inverted version of the mask (*inverted mask*) is used to produce the *non-photoconverted background* image that encompasses all the pixels in the image that are not segmented as part of the chromosome axes. The average intensity of the *non-photoconverted background* is then subtracted from the *masked non-photoconverted image* to generate the final *background subtracted non-photoconverted image*. *Bottom*. The *raw photoconverted image* signal outside the chromosome axes is eliminated using the *non-photoconverted mask* previously obtained. The photoconverted nuclei (outlined in red) are cropped out of the resulting image (*masked photoconverted image*) to get the *photoconverted channel background*. The average background intensity of the photoconverted channel was subtracted from the *masked photoconverted image* to obtain the final *background subtracted photoconverted image*. The last panel shows the merged photoconverted (green) and non-photoconverted (magenta) background subtracted merged images. The outlined nucleus corresponds to the example shown in Fig. S4B.

**C**, Steps used to obtain the photoconverted chromosome profile measurements. The images and plot profile correspond to a chromosome at time point *t=0 min.* after photoconversion. The photoconverted (green) and non-photoconverted (magenta) signals are shown with the crossover position indicated by the yellow arrowhead. The photoconverted chromosome trace is shown in blue. The trace is used to straighten the photoconverted chromosome. A plot of the average intensity of all the vertical pixels at every horizontal pixel position is generated and normalized to the mean of the values with the Y axis representing the normalized average intensity and the X axis representing the length of the photoconverted chromosome. The non-photoconverted mMaple::SYP-3 plot is shown in magenta and the photoconverted mMaple::SYP-3 plot is shown in green. The yellow rectangle represents the crossover region.

### Figure S5

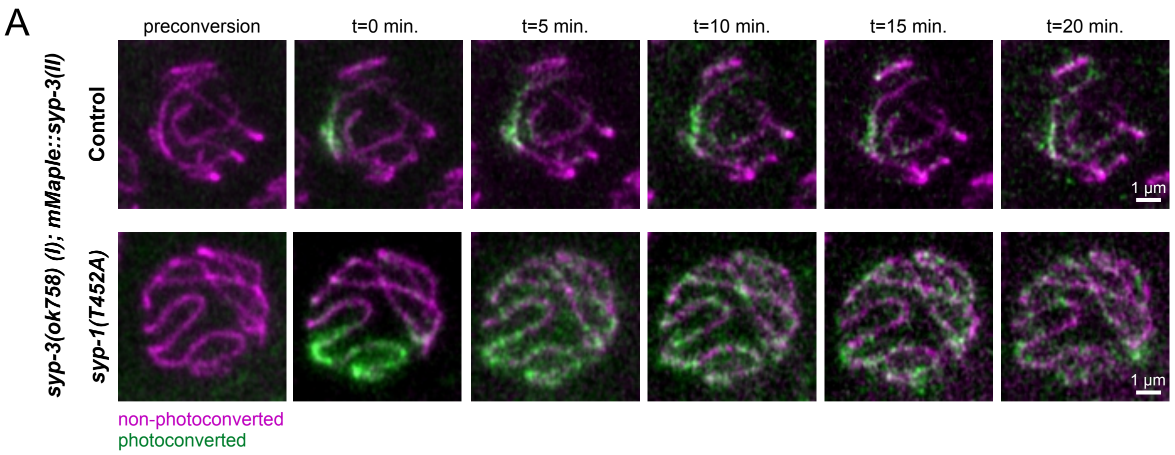
A
preconversion
t=0 min.
t=5 min.
t=10 min.
t=15 min.
t=20 min.
syp-3(ok758) (I); mMaple::syp-3(II)
Control
syp-1(T452A)
1 μm
1 μm
non-photoconverted
photoconverted

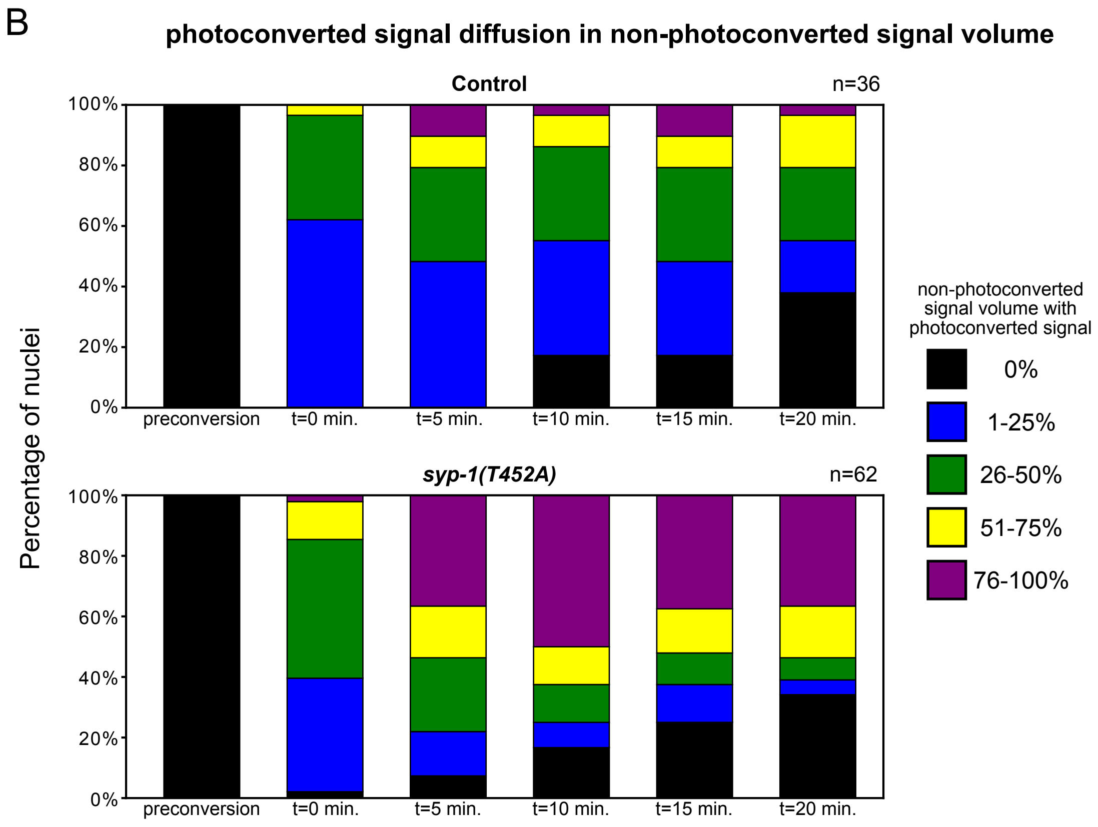
B
photoconverted signal diffusion in non-photoconverted signal volume
Control
n=36
syp-1(T452A)
n=62
Percentage of nuclei
100%
80%
60%
40%
20%
0%
preconversion
t=0 min.
t=5 min.
t=10 min.
t=15 min.
t=20 min.
non-photoconverted signal volume with photoconverted signal
0%
1-25%
26-50%
51-75%
76-100%

**Figure S5**. *Photoconverted mMaple3::SYP-3 diffuses faster and to a greater extent in syp-1(T452A) mutants*. Related to Figure 5.

**A**, Time course of *syp-3(ok758) (I); mMaple3::syp-3 (II)* (Control) and *syp-3(ok758) (I); mMaple3::syp-3 (II); syp-1(T452A) (V)* ( *syp-1(T452A)* ) showing the diffusion of photoconverted mMaple3::SYP-3 (green) throughout the non-photoconverted mMaple3::SYP-3 (magenta). The pre-conversion image is shown followed by 5 time points in 5 minute intervals from *t=0 min.* to *t=20 min*.

**B**, Stacked bar plots showing the ratio of nuclei classified within 5 categories (zero signal and four 25% quartiles) for the nonphotoconverted signal volume where the photoconverted signal had diffused at every time point. *Top.* Plot for the control nuclei (n=36). *Bottom.* Plot for the *syp-1(T452A)* nuclei (n=62). The categories for the volume occupied by the photoconverted signal are: 0% (black bars), 1-25% (blue bars), 26-50% (green bars), 51-75% (yellow bars) and 76-100% (purple bars).

### Figure S6 Partitioning simulation on various configurations

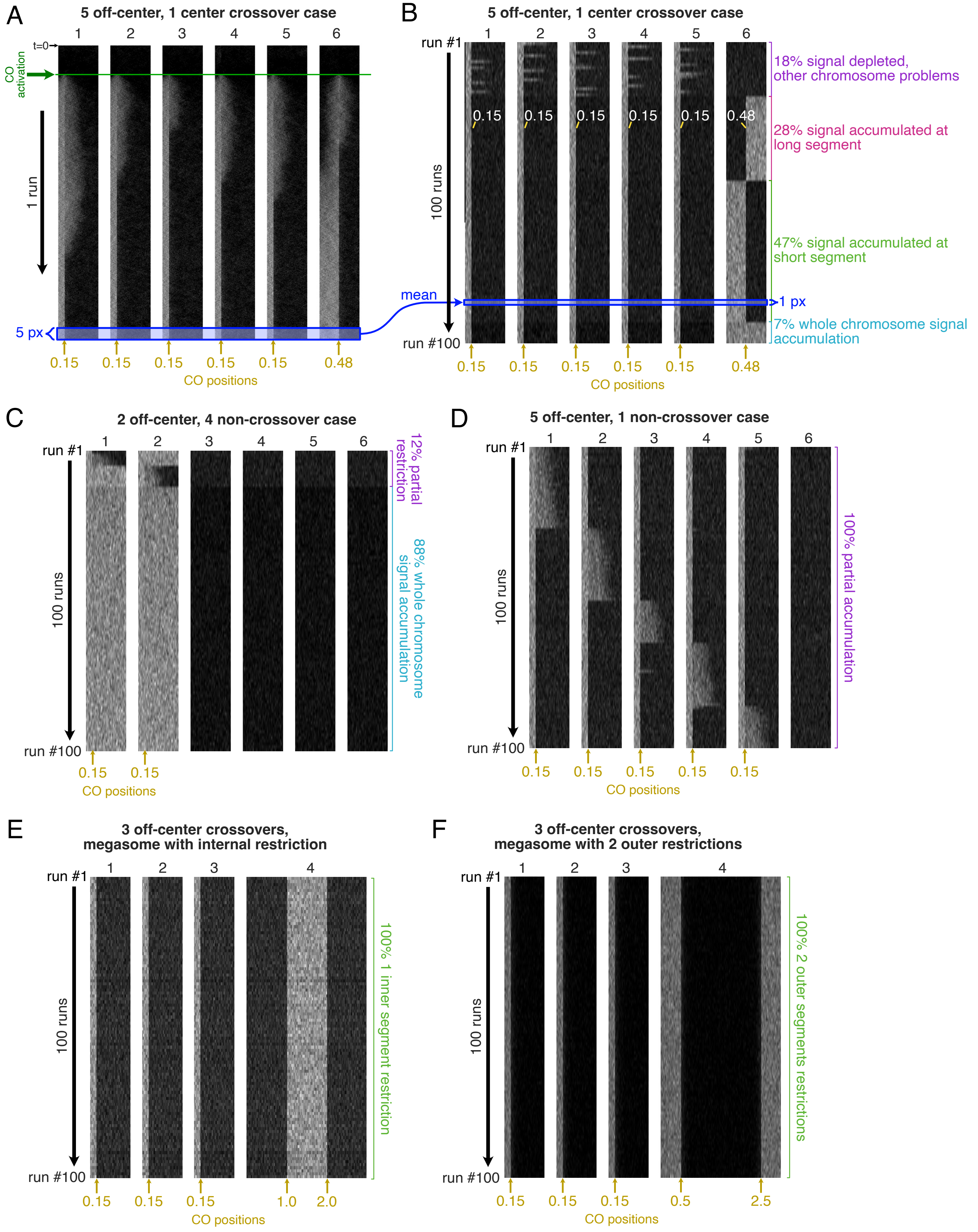

**Figure S6**. *Summaries of partitioning simulations on various configurations*. Related to Figure 6.

**A**, Output of 1 run of the simulation showing the spread of the signal along 6 chromosomes. All chromosomes have an arbitrary length of 1 (divided into 100 bins). The run consists of 15000 iterations. The state of the signal along each chromosome is represented as a row of pixels sorted vertically starting at t=0 at the top of the image. The horizontal pixels represent the amount of signal at that specific position in the corresponding chromosome. The crossovers are activated at t=1500 (green line) and their position is indicated by the yellow arrows and numbers at the bottom of the images. 5 chromosomes have off-center crossovers set at 0.15 and 1 chromosome has a near-center crossover set at 0.48. The blue box at the end of the run illustrates the last 5 states (box not to scale with the actual image) of each chromosome (see legend for **B-F**).

**B-F**, Summaries of 100 simulation runs with different crossover positions and chromosome configurations. Each vertical pixel row is the average intensity projection of the last 5 states of an individual run represented by the blue boxes and arrow shared between **Fig. S6A** and **Fig. S6B**. The crossover positions are marked with the yellow arrow and numbers. Normal chromosome lengths were set to 1 while megasome lengths were set to 3.

**B**, Summary of 100 simulation runs with 5 off-center crossover chromosomes (chromosomes #1-5) and 1 near-center crossover chromosome (chromosome #6). All off-center crossovers were set to position 0.15 and the near-center crossover was set to 0.48. The runs are sorted by the outcome of chromosome #6. Descriptions of each outcome class are at right: the purple range indicates the 18% of the runs where chromosome #6 was completely depleted of the signal; the pink range indicates runs in which chromosome #6 accumulated the signal on its long segment; the green range indicates where chromosome #6 accumulated the signal on its short segment; the light blue range indicates where chromosome #6 accumulated the signal at its entire length.

**C**, Summary of 100 simulation runs with 2 off-center crossover chromosomes and 4 non-crossover chromosomes. The off-center crossovers were set to position 0.15. The runs were sorted by the outcome of the off-center crossover chromosomes. The purple range indicates the 12% of runs where the off-center crossover chromosomes had partial restriction of the signal and the light blue range indicates the 88% of runs where the off-center chromosomes accumulated the signal along their entire length.

**D**, Summary of 100 simulation runs with 5 off-center crossover chromosomes and 1 non-crossover chromosome. The off-center crossovers were set to position 0.15. The runs were sorted by the outcome of the off-center crossover chromosomes. The purple range indicates that 100% of the runs had partial signal accumulation in at least one of the off-center chromosomes.

**E**, Summary of 100 simulation runs with 3 normal length chromosomes with off-center crossovers (at position 0.15) and 1 megasome with 2 near-center crossovers (at positions 1.0 and 2.0). The green range indicates that all simulated megasomes went on to produce signal restriction at a single inner segment.
**F**, Summary of 100 simulation runs with 3 normal length chromosomes with off-center crossovers and 1 megasome with 2 off-center crossovers. The crossover positions on the normal length chromosomes were set to 0.15. The crossover positions on the megasome were set to 0.5 and 2.5. The green range indicates that all simulated megasomes developed two outer segment restrictions.

| REAGENT or RESOURCE | SOURCE | IDENTIFIER |
|---|---|---|
| **Antibodies** | | |
| guinea pig-anti-SYP-1 | Carlton Lab | N/A |
| rabbit-anti-SYP-1phos | Carlton Lab | N/A |
| mouse-anti-GFP | Roche | Cat#12600500 |
| goat-anti-SYP-1 | Dernburg Lab | N/A |
| mouse-anti-HA | BioLegend | RRID:AB_2565006; Cat#901501 |
| mouse-anti-FLAG | SIGMA | RRID:AB_262044; Cat#F1804 |
| **Chemicals, peptides, and recombinant proteins** | | |
| Leibowitz L-15 without phenol red, 9.3% | gibco | Cat#21083027 |
| Cas9 | IDT | N/A |
| **Experimental models: Organisms/strains** | | |
| *C. elegans: GFP::cosa-1(II): meIs8 [pie-1p::GFP::cosa-1 + unc-119(+)] II* | Caenorhabditis Genetics Center | RRID:WB-STRAIN:WBStrain00000296; AV630 |
| *C. elegans: meT7 (III;X;IV): dpy-18(e364) unc-3(e151) meT7 III;X;IV.* | Caenorhabditis Genetics Center | RRID:WB-STRAIN:WBStrain00000292; AV311 |
| *C. elegans: GFP::cosa-1 (II); meT7 (III;X;IV): GFP::cosa-1(II): meIs8 [pie-1p::GFP::cosa-1 + unc-119(+)] II; dpy-18(e364) unc-3(e151) meT7 III;X;IV.* | This study | PMC468 |
| *C. elegans: syp-1(T452A)::HA (II); syp-1(me17) (V); syp-1(icm81[T452A::HA]) II; syp-1(me17)/nT1[unc-?(n754) let-? qIs50] IV;V* | This study | PMC471 |
| *C. elegans: GFP::cosa-1 (II); syp-1(T452A)::HA (V): meIs8 [pie-1p::GFP::cosa-1 + unc-119(+)] II; syp-1(icm85[T452A]) /nT1[qIs 51] IV; V* | This study | PMC703 |
| *C. elegans: syp-1(T452A); icmSi44 [Psyp-1::syp-1(T452A) + unc-119(+)] II* | Sato-Carlton et al[19] | PMC346 |
| *C. elegans: dtn-1::3xFLAG (II): dtn-1::3FLAG::GFP II* | Yamamoto et al[34] | PHX2016 |
| *C. elegans: dtn-1::3xFLAG (II); meT7 (III;X;IV): dtn-1::3Flag::GFP II; meT7 [dpy-18(e364) unc-3(e151)] III;X;IV* | This study | PMC716 |

| | | |
|---|---|---|
| *C. elegans: syp-3(ok758) (I); GFP::cosa-1 mMaple3::SYP-3 (II): syp-3(ok758) I; ieSi63 [cbunc-119+, psyp-3::mMaple3::syp-3] II; unc-119(ed3) III.* | Rog et al[9] | CA1234 |
| *C. elegans: syp-3(ok758) (I); mMaple3::syp-3 (II); syp-1(T452A) (V): syp-3(ok758) I; ieSi63 [cbunc-119+, psyp-3::mMaple3::syp-3] II; unc-119(ed3) III; syp-1(icm85[T452A]) V/nT1[qls 51] IV; V* | This study | PMC715 |
| **Oligonucleotides** | | |
| Primer: c02_t452a_ha_4_fw: (5'-ACCCATACGACGTCCCAGACTA-3') | This study | N/A |
| Primer: c02_t452a_ha_4_rev: (5'-CGCTAAAAACACTAATCACACGGA-3') | This study | N/A |
| Primer: c02_t452a_ty1_fw: (5'-TTCAGATCGAGTAGTTCCGCCG-3') | This study | N/A |
| crRNA: c02_t452a_ha_crRNA2: (5'-GGAAGAAATAATGTGTGTGT-3') | This study | N/A |
| Ultramer: c02_t452a-ha_hrt_ultramer: (5'-GAGAGCCGAAGCTCATACTGCAGATGTTCGCCGAAAGAGAGGAGGGAAGAAAGAGGTCCACACCAACCAGGACCCACTCGACTACCCATACGACGTCCCAGACTACGCTTAATGTGTGTGTGCCGAAGAAACGACTATGTACCATTTCAATCTTGTGCTATTTTTTTTTGTGTTTTTAGAGTTTTATATAAAGTAATTTG-3') | This study | N/A |
| **Software and algorithms** | | |
| Fiji v2.14.0 | Schindelin et al[51] | RRID:SCR_002285; https://imagej.net/software/fiji/ |
| Priism | Chen et al[52] | N/A |
| ZEN (Black edition) | ZEISS | RRID:SCR_018163 |
| tifffile v2022.5.4 | Gohlke[53] | RRID:SCR_023338; https://zenodo.org/records/6795861 |
| roifile v2023.8.30 | Gohlke[55] | RRID:SCR_023331; https://zenodo.org/records/8303336 |
| Python 3.12 | Python Software Foundation | RRID:SCR_008394; https://www.python.org/ |

| | | |
|---|---|---|
| SNT framework v4.2.1 | Arshadi et al[54] | RRID:SCR_016566; https://imagej.net/update-sites/neuroanatomy/ |
| Trainable Weka Segmentation v3.3.4 | Arganda-Carreras et al[57] | https://imagej.net/plugins/tws/ |
| SciPy v1.11.2 | Virtanen et al[58] | RRID:SCR_008058; https://scipy.org/ |
| **Other** | | |
| All original code used in this study | This study | https://www.github.com/carltonlab/chromosome-partitioning |
